# Superiority of chromosomal compared to plasmid-encoded compensatory mutations

**DOI:** 10.1101/2024.01.15.575717

**Authors:** Rosanna C.T. Wright, A. Jamie Wood, Michael J. Bottery, Katie J. Muddiman, Steve Paterson, Ellie Harrison, Michael A. Brockhurst, James P.J. Hall

**Affiliations:** Division of Evolution, Infection and Genomic Sciences, University of Manchester, Manchester, United Kingdom; Department of Biology, University of York, York, United Kingdom; Department of Mathematics, University of York, York, United Kingdom; Department of Evolution, Ecology and Behaviour, Institute of Infection, Veterinary and Ecological Sciences, University of Liverpool, Liverpool, United Kingdom; School of Biosciences, University of Sheffield, Sheffield, United Kingdom

## Abstract

Plasmids are important vectors of horizontal gene transfer in microbial communities but can impose a burden on the bacteria that carry them. Such plasmid fitness costs are thought to arise principally from conflicts between chromosomal- and plasmid-encoded molecular machineries, and thus can be ameliorated by compensatory mutations (CMs) that reduce or resolve the underlying causes. CMs can arise on plasmids (i.e. plaCM) or on chromosomes (i.e. chrCM), with contrasting predicted effects upon plasmid success and subsequent gene transfer because plaCM can also reduce fitness costs in plasmid recipients, whereas chrCM can potentially ameliorate multiple distinct plasmids. Here, we develop theory and a novel experimental system to directly compare the ecological effects of plaCM and chrCM that arose during evolution experiments between *Pseudomonas fluorescens* SBW25 and its sympatric mercury resistance megaplasmid pQBR57. We show that while plaCM was predicted to succeed under a broader range of parameters in mathematical models, experimentally chrCM dominated under all conditions, including those with numerous recipients, due to a more efficacious mechanism of compensation, and advantages arising from transmission of costly plasmids to competitors (plasmid ‘weaponisation’). We show analytically the presence of a mixed Rock-Paper-Scissors regime for plaCM, driven by trade-offs with horizontal transmission, that explains the observed failure of plaCM to dominate even in competition against an uncompensated plasmid. Our results reveal broader implications of plasmid-bacterial evolution for plasmid ecology, demonstrating the importance of compensatory mutations for resistance gene spread. One consequence of the superiority of chrCM over plaCM is the likely emergence in microbial communities of compensated bacteria that can act as ‘hubs’ for plasmid accumulation and dissemination.

## Background

Conjugative plasmids are important for bacterial evolution. Plasmids transfer niche-adaptive ecological functions between lineages and consequently can drive adaptation and genomic divergence (Finks and Martiny, 2023; Vos et al., 2023; Wein and Dagan, 2020). However, acquiring a new conjugative plasmid is frequently costly for the host cell. Such plasmid fitness costs can arise from a variety of causes, including the metabolic burden of plasmid maintenance, disrupted gene regulation, stress responses, cytotoxicity, and mismatched codon usage (San Millan and MacLean, 2017). The long-term persistence of costly plasmids within bacterial lineages often requires compensatory evolution to negate these fitness costs (Brockhurst and Harrison, 2022). Experimental evolutionary studies have revealed that compensatory mutations (CMs) may occur on the plasmid, or the chromosome, or both replicons (Benz and Hall, 2022; Bottery et al., 2017; Dahlberg and Chao, 2003; De Gelder et al., 2007; Hall et al., 2021, 2019; Harrison et al., 2015; Jordt et al., 2020; Loftie-Eaton et al., 2017; San Millan et al., 2015; Stalder et al., 2017). Such CMs affect a wide range of gene functions, including regulatory genes, helicases, other co-resident mobile genetic elements, or hypothetical genes without known function.

The fitness cost of a given plasmid in a given host can be ameliorated by alternative CMs affecting distinct genetic targets, sometimes encoded by different replicons, i.e., affecting genes on the chromosome, which we term chrCM, or on the plasmid, which we term plaCM. This phenomenon is exemplified by the common soil bacterium *Pseudomonas fluorescens* SBW25 (henceforth ‘SBW25’) and the environmental mercury resistance plasmid pQBR57. Both SBW25 and pQBR57 were isolated from sugar beet plants at a field site in Wytham Woods, Oxfordshire, UK in the 1990s (Bailey et al., 1995; Lilley et al., 1996). pQBR57 causes a substantial fitness cost in SBW25 due to a specific genetic conflict with a chromosomal hypothetical gene PFLU4242, inducing a sustained SOS response and the maladaptive expression of chromosomal prophages leading to cell damage (Hall et al., 2021). This costly cellular disruption can be negated by single CMs affecting either PFLU4242 itself or a plasmid-encoded regulator PQBR57_0059. Either CM is sufficient to reduce the fitness cost of pQBR57 and both fix the transcriptional disruption caused by pQBR57 acquisition. For clarity, we refer to SBW25 strains with a loss-of-function mutation in PFLU4242 as SBW25::chrCM to indicate chromosomal CM, and strains with a loss-of-function mutation in PQBR57_0059 as pQBR57::plaCM to indicate plasmid CM. Although both CMs evolved in SBW25 plasmid-carrying populations in potting soil microcosms, they were never observed to co-occur in the same genome, suggesting that there is no added benefit of combining both CMs in the same cell. chrCM can ameliorate the fitness costs of other pQBR plasmids, whereas the benefits of plaCM are transmitted when pQBR57::plaCM transfers by conjugation (Hall et al., 2021, 2019).

Where alternative CMs exist on the chromosome or the plasmid, existing theory predicts that plaCMs will be superior (Zwanzig et al., 2019). This superiority arises because, unlike chrCMs which are only inherited vertically at cell division, plaCMs are also transmitted horizontally by conjugation. Provided the plaCM also negates the fitness cost of the plasmid in newly formed transconjugant cells, the linkage of the plasmid and the CM can thus enhance plasmid maintenance and spread. Correspondingly, plaCMs are predicted to outcompete chrCMs even if plaCMs offer less efficient amelioration than chrCMs. However, previous attempts to explore these predictions theoretically have relied on numerical simulations and extensive parameter fitting (Rebelo et al., 2023a; Zwanzig et al., 2019), limiting the generalisability of the findings, while experimental tests competing alternate modes of CM are lacking altogether. Moreover, some experiments have reported a trade-off between CMs and conjugation rate, which could impede the success of plaCMs (Bethke et al., 2023; Dimitriu et al., 2021; Turner et al., 1998).

To contrast the effect of chrCM with plaCM on bacteria-plasmid dynamics, we first develop two simple mathematical models based on 3-equation ordinary differential equations (ODEs) in which we consider the arrival of a plasmid-bearing strain with either a chrCM or a plaCM. A key strength of this approach is that the models we create can be solved exactly to provide general understanding. The predictions generated by our models were then tested experimentally. To enable direct competition of chrCM and plaCM we engineered variants of SBW25 and pQBR57 carrying defined CMs and fluorescent tags allowing cells containing the plaCM and/or chrCM to be distinguished by flow cytometry. We then performed competition experiments across various ecological scenarios predicted to alter the differential benefits of these contrasting modes of compensatory evolution. Specifically, we varied the strength of mercury selection, the presence of other plasmid replicons in the population, or the availability of plasmid-free recipient cells for onward conjugative transfer within the population. We show that, contrary to expectations, plaCM performs poorly in competition with chrCM under all tested conditions due to lower efficacy of amelioration, a probable trade-off against conjugative efficacy, and an overlooked benefit of chrCM that effectively enables these cells to ‘weaponise’ costly plasmids to reduce the fitness of competitors. Our results have implications for the mobilisation of genes in microbial communities, suggesting that chromosomes are more likely to become plasmid-favourable — and thus hubs of source-sink horizontal gene transfer — than plasmids are to become low-cost generalists across hosts.

## Results

### Mathematical model of plasmid- or chromosome-encoded compensatory mutation dynamics

To understand the dynamics of chrCM and plaCM we modified a simple, well-understood model of bacteria-plasmid population dynamics to include either chrCM or plaCM. Separate models were preferred because the combined system (i.e., containing both chrCM and plaCM) is too complex to solve analytically. The basic model, without CMs, is detailed in the Supplementary Appendix and recapitulated the key findings of previous studies wherein costly plasmids do not invade unless their conjugation rate, 𝛾, is larger than 𝜇 (𝛼 − 𝛽)⁄(𝛼 − 𝜇), and only competitively displace the plasmid-free population if 𝛾 is larger than 𝜇 (𝛼 − 𝛽)⁄(𝛽 − 𝜇), where 𝛼 is the plasmid free growth rate, 𝛽 is the plasmid containing growth rate, and 𝜇 is the population turnover rate. Positive selection for the plasmid is included via the selection pressure term, 𝜂, which is initially set to zero.

We then considered how the addition of chrCM affects the outcome of this underlying basic system. Here, plasmid free wild-type bacteria (*f*) and wild-type bacteria containing a wild-type plasmid (*p*) are invaded by a chrCM variant bearing a wild-type plasmid (*c*). The compensatory effect is assumed to be imperfect so the growth rate of the three strains are assumed to be 𝛼 for *f*, 𝛽_𝑃_ for *p*, and 𝛽_𝐶_ for *c*, where 𝛼 > 𝛽_𝐶_ > 𝛽_𝑃_. The dynamics of the system are then described by the following set of ODEs:

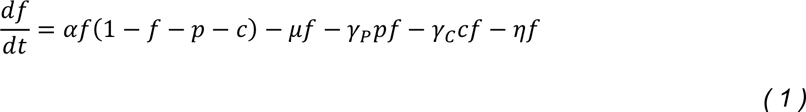

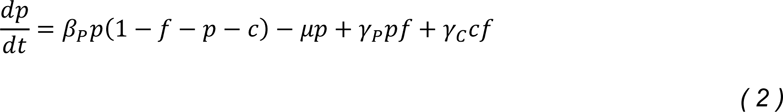

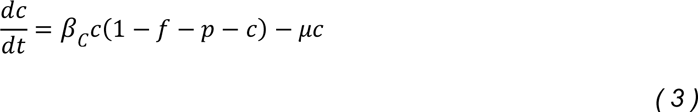

In the case of no selection for the plasmid, 𝜂 = 0, this set of ODEs can be solved exactly to yield a simple phase plane structure in which chrCM sweeps to fixation (0,0,*c*\*) from the expected stable fixed point of the underlying system according to the value of the conjugation rate (Fig. 1A). When the underlying fixed point is plasmid-free only (*f*\*,0,0) (i.e. 𝛾_𝑃_ < 𝜇 (𝛼 − 𝛽_𝑃_)⁄(𝛼 − 𝜇)) or mixed (*f*\*,*p*\*,0) (i.e. 𝜇 (𝛼 − 𝛽_𝑃_)⁄(𝛽_𝑃_ − 𝜇) > 𝛾_𝑃_ > 𝜇 (𝛼 − 𝛽_𝑃_)⁄(𝛼 − 𝜇)), and compensation is imperfect, the fixed points are separated by a saddle at (*f*s,*p*s,*c*s). This means that when the conjugation rate is sufficiently high the chrCM will always invade, but at lower conjugation rates (provided 𝛼 > 𝛽_𝐶_) there is a threshold. If the initial proportion of chrCM exceeds the threshold value (given by the saddle), complete replacement occurs and chrCM successfully invades and moves to fixation. The reason invasion can occur despite chrCM having lower growth than the plasmid-free is that a plasmid-carrying chrCM can conjugate the costly plasmid into plasmid-free competitors, such that chrCM then exceeds the uncompensated competitor’s growth rate (i.e. 𝛽_𝐶_ > 𝛽_𝑃_) and takes over the system. Selection (𝜂 > 0) reduces the relative ability of plasmid-free wild-type *f* to compete against plasmid-bearers, concentrating the dynamics on the competition between *p* and *c* and the difference between 𝛽_𝐶_ and 𝛽_𝑄_, thus favouring chrCM. This result is demonstrated in the Supplemental Appendix, and typically occurs with a linear addition to the stability conditions which rapidly favour the chrCM plasmid-bearing invader.

**Figure 1.**
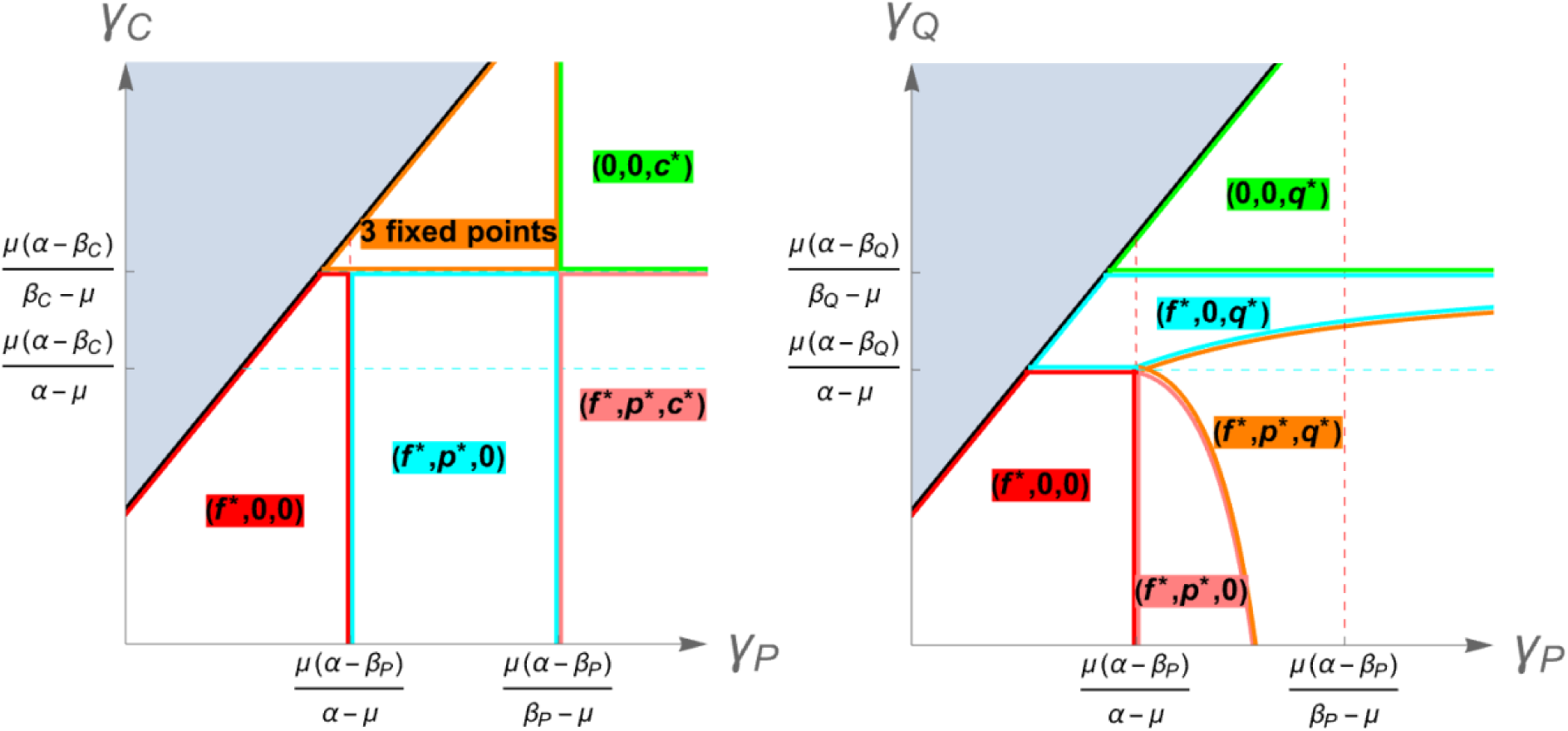
Phase portraits describing the fate of compensatory mutations. (A) Chromosomal CMs. (B) Plasmid CMs. Axes describe relative conjugation rates without (x) and with (y) the corresponding CM, with the black line indicating no difference. Where 𝛾_𝐶𝑀_ > 𝛾_𝑃_ (i.e. both growth rate and conjugation rate are increased by the CM, indicated in grey) the wild-type plasmid is always lost with the system reverting to a two-member basic model.

We next considered a system invaded by a variant bearing a plaCM plasmid (*q*). This leads to a set of three differential equations analogous to those above:

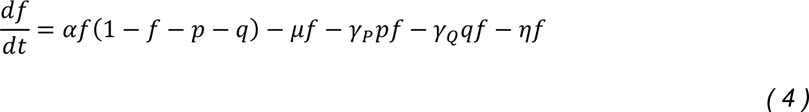

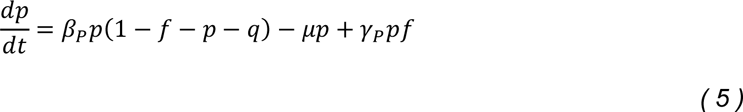

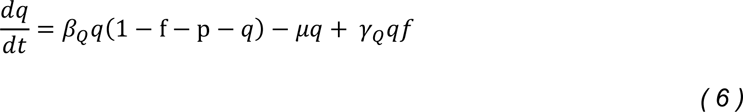

Analysis of the system reveals a more complex phase portrait, described in Fig. 1B in the case of no selection, 𝜂 = 0. Consistent with prior work (Zwanzig et al., 2019), comparison of Figs 1A and 1B shows a wider range of conditions in which plaCM is successful compared with chrCM. Where there is no trade-off with conjugation rate, plaCM always displaces the uncompensated plasmid, invading the system if 𝛾_𝑄_ < 𝜇 (𝛼 − 𝛽_𝑄_)⁄(𝛼 − 𝜇) and dominating if 𝛾_𝑄_ > 𝜇 (𝛼 − 𝛽_𝑄_)⁄(𝛽_𝑄_ − 𝜇), and, unlike chrCM, in a manner that does not depend on initial conditions.

Previous experiments have shown that CMs can affect the ability of a plasmid to conjugate (Bethke et al., 2023; Dimitriu et al., 2021; Turner et al., 1998). We therefore investigated how the success of each CM is affected by changes in the conjugation rate. For plaCM, if 𝛾_𝑄_ > 𝛾_𝑃_, i.e. plaCM confers a higher transfer rate than the wild-type plasmid, the wild-type plasmid is always lost from the system, and the outcome for plaCM collapses into the single-plasmid system described above (Supplementary Appendix). However, if there is a trade-off such that 𝛾_𝑃_ > 𝛾_𝑄_, various outcomes are possible depending on the other parameters, including loss of both plasmids (*f*\*,0,0), fixation of plaCM (0,0,*q*\*), co-existence between wild-type and plasmid-free (*f*\*,*p*\*,0), and, unexpectedly, a state with a stable coexistence between the *f*, *p* and *q* populations (Fig. 1B orange region), which would not be found in a linearised adaptive dynamics approach. The stable fixed point is oscillatory in character (a stable spiral) and is driven by Rock-Paper-Scissors (RPS)-like nontransitive dynamics. When *f* is large, this promotes the conjugative spread of the fastest conjugating population, *p*. When *p* is large, *f* is small, so the opportunities for conjugation are relatively low but the force of infection remains high due to high 𝛾_𝑃_. Here, the *q* outgrows the *p*. When *q* is large opportunities for conjugation are also low but as 𝛾_𝑄_ is relatively low the *f* can outgrow the *q*. This RPS-like dynamic is approximate as the interactions are not perfectly symmetric: in the prototypical model each type has a direct impact on one other type and is directly impacted by the third. Here the competition arises though a mixture of competitive growth and competitive infection and is different for each combination of subpopulations.

For chrCM, the system is robust to changes in conjugation rate provided compensation is sufficiently strong (𝛽_𝐶_ > 𝜇(𝛾_𝐶_ + 𝛼)/(𝜇 + 𝛾_𝐶_)). In cases where chrCM has a more substantial effect on 𝛾_𝐶_, the CM is lost (Fig. 1A red and blue regions), except in cases where the conjugation rate from uncompensated strains is sufficiently high (Fig. 1A pink area, 𝛾_𝑃_ > 𝜇(𝛼 − 𝛽_𝑃_)/(𝛽_𝑃_ − 𝜇)). Under these conditions, the force of infection of costly plasmids ensures that the frequency of plasmid-bearers in the system is maintained at a high enough level such that chrCM has sufficient competitive advantage to persist. Although the chrCM system can also admit an oscillatory stable coexistent solution, driven by a form of RPS dynamics it is only for 𝛾_𝑃_ large, 𝛾_𝐶_ small and further away from the biologically relevant regime.

Selection simplifies the dynamics of the plaCM system by reducing the effect of the plasmid-free population (*f*) to the dynamics. Even when 𝛾_𝑃_ ≫ 𝛾_𝑄_, plaCM is more likely to invade across a range of parameters because the contribution of the conjugation terms to the dynamic is reduced, resulting in a head-to-head competition between *p* and *q*, which the latter will always dominate due to lack of infection opportunities which would be granted by a large plasmid free *f* population.

Although four-equation models describing a direct competition between chrCM and plaCM cannot be solved analytically, some insight can be gained by comparing equations 3 and 6. Specifically, we can see that as the abundance of plasmid-free recipients decreases, the 𝛾_𝑄_𝑓 component that positively affects the success of plaCM correspondingly decreases, such that the relative success of plaCM and chrCM is increasingly determined by the difference between 𝛽_𝐶_ and 𝛽_𝑞_, i.e. the relative strengths of amelioration of the two CMs.

Overall, then, our models identify a broader range of parameter space in which plaCM is likely to succeed, relative to chrCM. However, we also predict that trade-offs between compensatory mutations and plasmid conjugation can have complex effects on the overall success of a CM, particularly in communities that contain a mixture of uncompensated plasmid-carrying and plasmid-free competitors. Selection reduces the complexity of the dynamics in both cases by removing potential recipients from the system, thus reducing the contribution of plasmid transmission to the dynamics, and increasing the dependency of the outcome on the relative strength of the CM.

### Knock-outs of putative ‘cost’ genes recapitulate compensatory mutations

To experimentally investigate the dynamics of plaCM *versus* chrCM, we established an experimental system in which fluorescently-labelled strains were engineered with plaCM or chrCM, enabling enumeration by flow cytometry. To test that our newly-engineered strains exhibited the fitness effects by flow cytometry that we have previously observed by CFU plating, we first performed 24-hour competition experiments. As expected, each CM ameliorated the fitness cost of plasmid carriage (Fig. 2, linear model [LM] one-sided posthoc comparison against 0: plaCM t_9_ = 22.1, p < 1e-7; chrCM t_9_ = 23.5, p < 1e-7). In addition, our head-to-head experimental design revealed that chrCM was marginally fitter than plaCM (coefficient = 0.13, t_9_ = 2.75, p = 0.045) and there was no detected fitness benefit of combining both CMs in the same cell (coefficient comparing single vs. double-compensation = −0.1, t_9_ = −1.86, p = 0.19). Interestingly, the benefits of CMs relative to non-compensated plasmid-carrying strains were enhanced in the presence of mercury, and more strongly so for chrCM than for plaCM (chrCM difference in coefficients = 1.0, plaCM difference in coefficients = 0.7), suggesting that the costly cellular disruption caused by pQBR57 is further exacerbated by exposure to mercury (Carrilero et al., 2021). These results cause us to expect acceleration of CM invasion under mercury selection.

**Figure 2.**
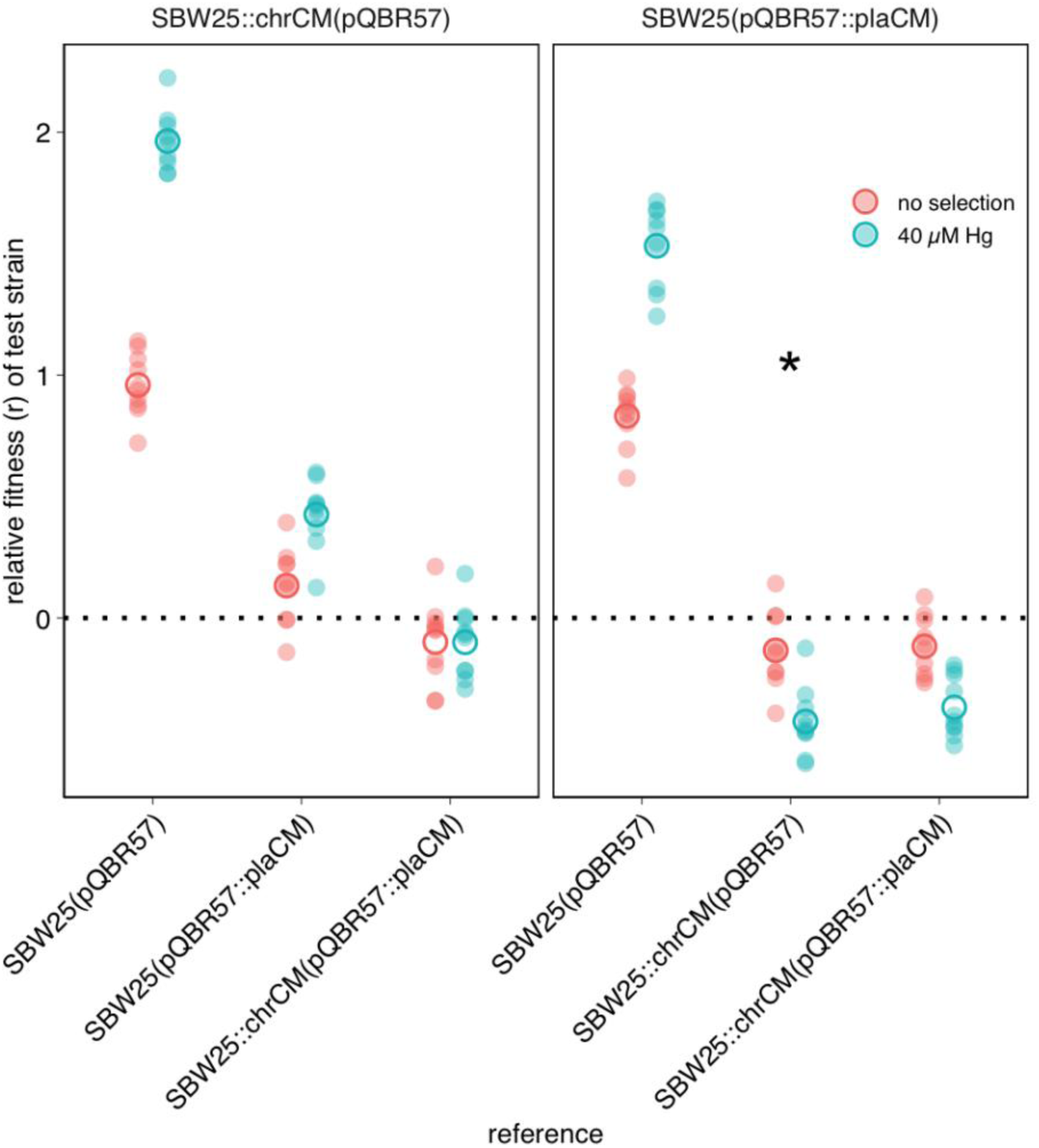
Engineered plaCM and chrCM variants both ameliorate plasmid fitness costs, though chrCM is more effective. Panels indicate test strains. Unfilled circles indicate mean across 10 replicates, each of which is indicated by a semi-transparent filled circle. The asterisk indicates data in the right panel which is also presented in the left panel.

### Chromosomal CM outcompetes plasmid-borne CM across various ecological conditions

To test our predictions, we used our validated strains to test the relative success of chrCM *versus* plaCM under varying ecological conditions. First, we varied the strength of selection for the plasmid-encoded mercury resistance trait. Informed by our models, we predicted that selection would increase the success of chrCM relative to plaCM. This is because selection would remove plasmid-free recipients from the system, and as plasmids prevent superinfection by similar plasmids through surface exclusion and/or entry exclusion, plaCM would gain no benefit from its ability to transfer by conjugation. For these experiments, our strains were labelled to track the relative success of each mode of compensation, and so the label was inserted into the replicon encoding the compensatory mutation i.e. the SBW25ΔPFLU4242 chromosome for chrCM, and pQBR57ΔPQBR57_0059 for plaCM. SBW25::chrCM carrying wild-type pQBR57 was competed against pQBR57::plaCM in a wild-type chromosomal background in populations initially containing wild-type plasmid-free recipients at 50% frequency (1:1:2 chrCM:plaCM:recipients), and populations were transferred for 8 transfers (∼50 generations). Consistent with our predictions, mercury selection indeed favoured chrCM (Fig. 3, Generalised Linear Mixed Effects Model [GLMM] interaction effect of selection:transfer 𝜒^2^ = 102.9, p < 1e-7, Fig. 3). However, chrCM was also fitter than plaCM without selection (coefficient for transfer = −0.32, z = −15.7, p < 2e-16), suggesting that any benefits from the ability of the plaCM to transfer into the recipient pool were outweighed by the reduced amelioration provided by plaCM relative to chrCM (Fig. 3).

**Figure 3.**
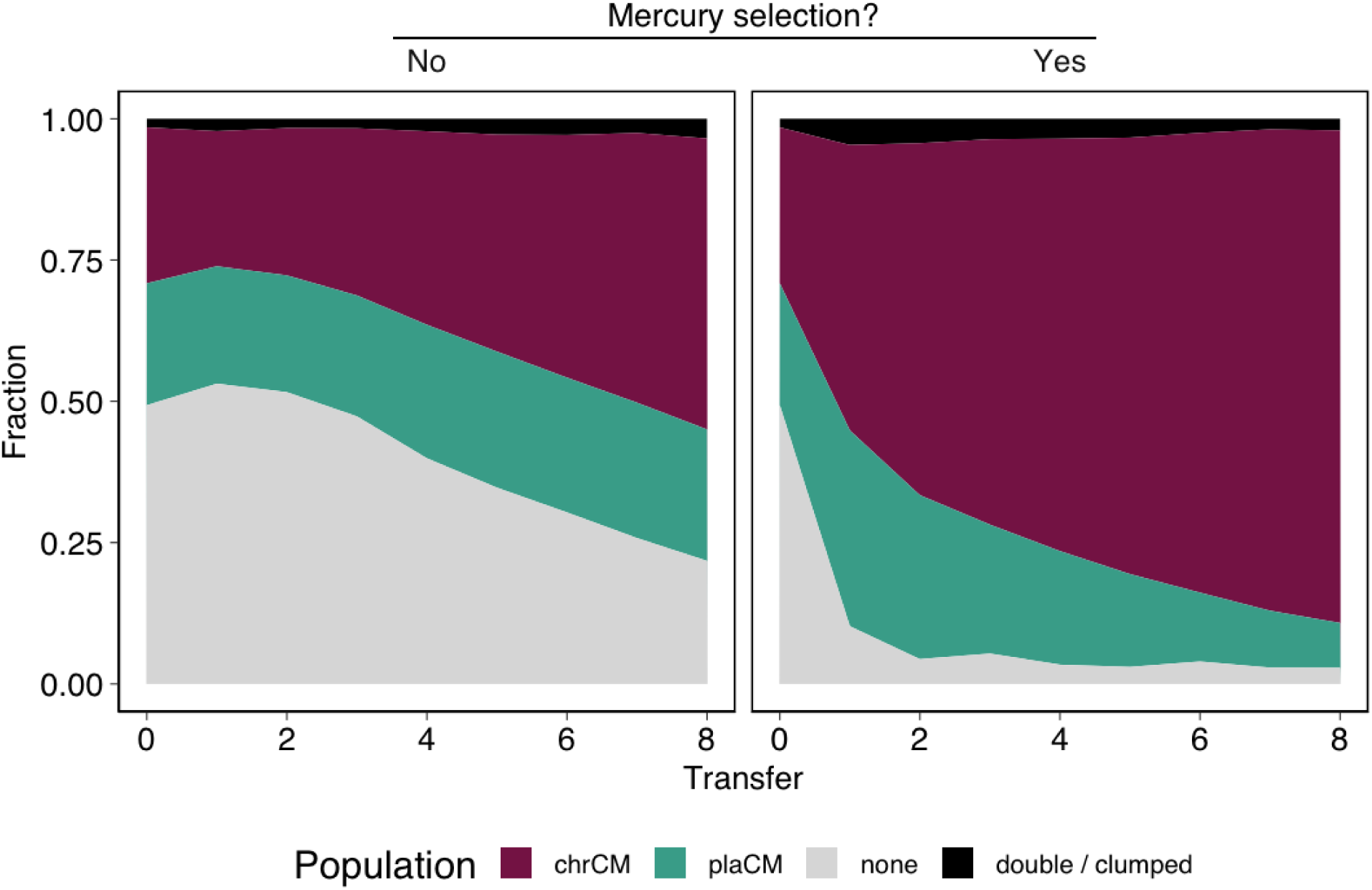
Chromosomal compensatory mutations outperform plasmid-borne compensatory mutations, particularly under positive selection. Subpanels indicate the presence of mercury selection (No/Yes). Mean of 20 replicates/treatment (10 per marker orientation). Individual replicate plots are shown in Figure S1.

In parallel, we investigated whether the benefits of chrCM could be increased in microbial communities hosting multiple costly plasmids. Our previous worked showed that pQBR57 and pQBR103 could be harboured in the same cell. We also showed pQBR103 could be compensated by the chrCM, ΔPFLU4242, both by itself and together with pQBR57, whereas evidence suggested that plaCM exacerbated the cost of pQBR103 (Carrilero et al., 2021; Hall et al., 2021). We therefore predicted that chrCM would be favoured over plaCM both with and without selection in pQBR103-harbouring communities, since chrCM would ameliorate both plasmids. Indeed, in communities harbouring pQBR103, chrCM again outcompeted plaCM (Fig. 4, GLMM coefficient for transfer = −0.30, z = −14.5, p < 2e-16, Fig 4), but there was no detectable additional effect of pQBR103 on the relative success of chrCM *versus* plaCM.

**Figure 4.**
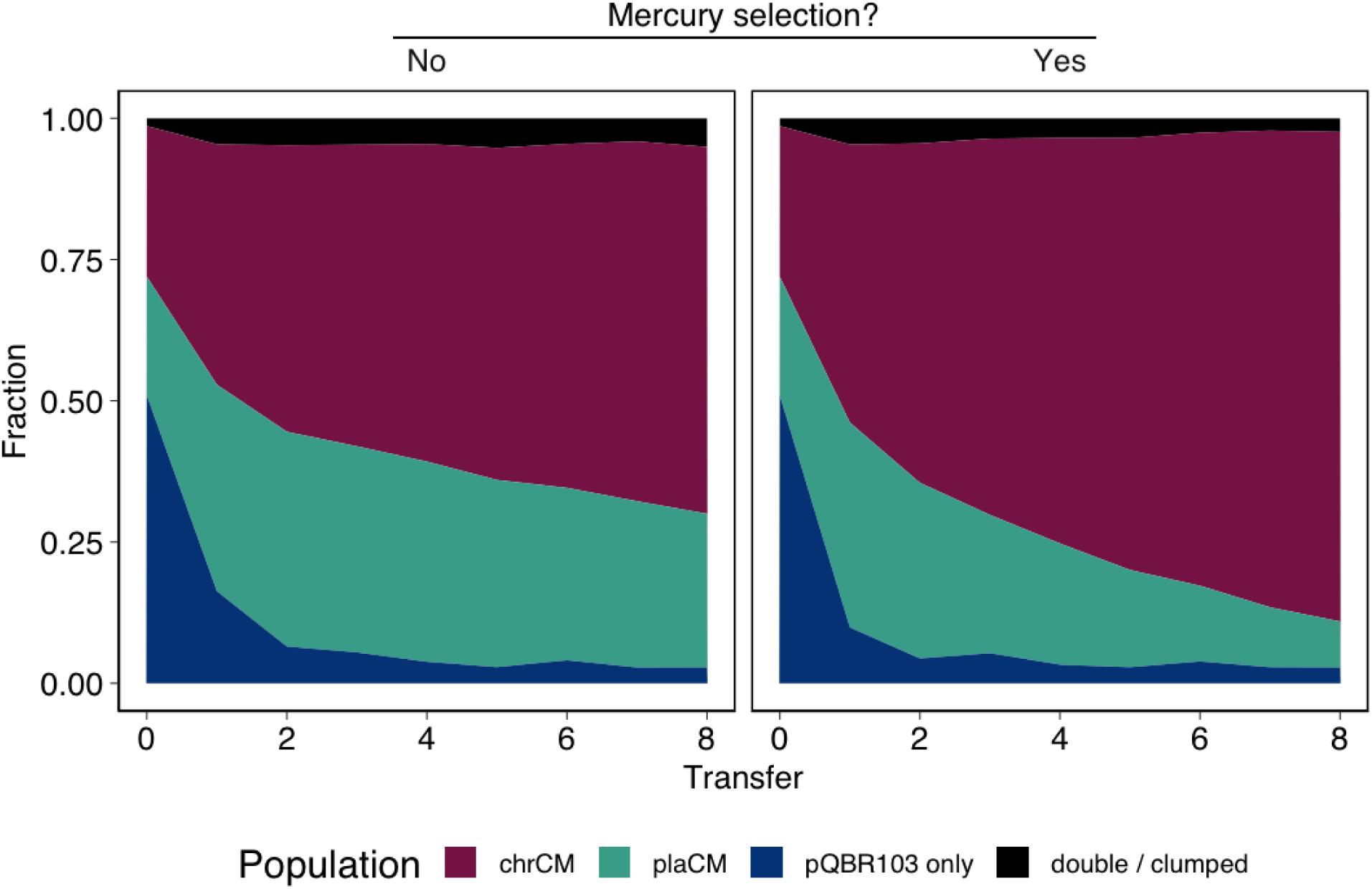
Chromosomal compensatory mutations were not additionally favoured in environments with another costly plasmid. Subpanels indicate the presence of mercury selection (No/Yes). Mean of 20 replicates/treatment (10 per marker orientation). Individual replicate plots are shown in Figure S2.

We reasoned that increasing the pool of potential recipients may tip the balance towards plaCM, since conjugation (and thus transmission of the CM) could then play a bigger role in the population dynamics. We again established populations beginning with equal proportions of plaCM and chrCM but varied the proportions of recipients: (i) 10-fold excess of plasmid-free; (ii) equal abundance of plasmid-free; (iii) 10% plasmid-free; (iv) without added plasmid-free. Populations were propagated without selection. Contrary to expectations, plaCM performed relatively poorly against chrCM across all frequencies of plasmid-free recipients (Fig. 5, GLMM transfer:ratio interaction 𝜒^2^ = 5.00, p = 0.17; main effect of transfer 𝜒^2^ = 88.6, p < 1e-7; coefficient for transfer = −0.35, z = −30.8, p < 2e-16), suggesting that the potential benefits of CM transmission could not be manifested by the plaCM.

**Figure 5.**
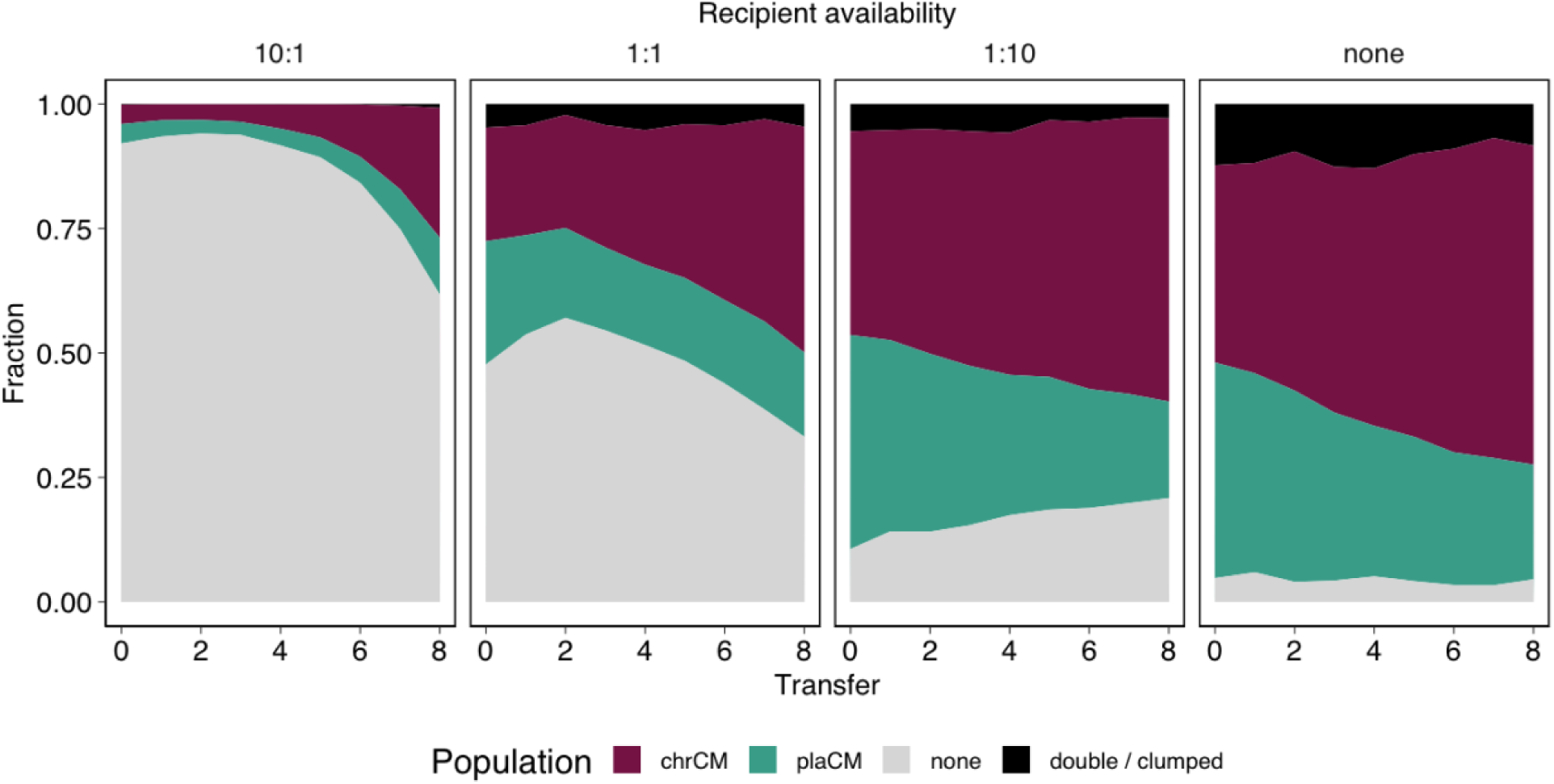
ChrCM was more successful than plaCM regardless of recipient availability. Sub-panels indicate, from left to right, the starting fraction of wild-type/plasmid-free cells relative to a 50:50 mix of chromosomal CM (with wild-type plasmid) and plasmid CM in wild-type cells. Plots indicate the mean of 6 independent experiments; replicate-level plots are provided in Figure S3.

Our model predicted that transmission of wild-type plasmids from chrCM cells could play an important role in the success of chrCM. Furthermore, the theoretical advantage gained by plaCM through horizontal transfer may be reduced when in competition against plasmids carried by cells with chrCM, because wild-type plasmid transfer from chrCM could remove potential recipients for plaCM. However, our initial experiments used fluorescent labels to track the fates of the different compensatory alleles, i.e. the chromosomes of chrCM (SBW25::chrCM), and the plasmids of plaCM (pQBR57::plaCM), rather than the plasmids which began in each background. To understand how the plasmids themselves were affected by the different CMs, we established a complementary experiment to that in Fig. 5, except, rather than tracking chrCM, we tracked the wild-type pQBR57 that began in the chrCM background (Fig. 6). Compared with pQBR57::plaCM, uncompensated pQBR57 from chrCM was significantly more successful, with a considerable proportion of the plasmids at the end of the experiment being uncompensated plasmids that began in the chrCM population, and patterns of invasion depending on the abundance of potential recipients (GLMM third-order polynomial transfer:ratio interaction 𝜒^2^ = 287.4, p < 1e-7). Notably, comparison with the experiments in Fig. 3 (which were performed in parallel) revealed that invasion of the plasmid from chrCM pre-empted the invasion of chrCM, indicating that plasmid transmission likely contributed to the overall success of chrCM. Essentially, as plasmid-free competitors are removed from the system by infection with the wild-type uncompensated plasmid, the space is filled by competition between CMs, which favours the more effective mechanism of compensation, chrCM. Indeed, when plasmid-free recipients were initially rare, the dynamics essentially come down to a competition between the compensatory mutations, with the most effective mechanism, chrCM, becoming dominant. The experimental results were therefore consistent with our model prediction that transmission of costly plasmids, and the consequent removal of plasmid-free recipients, facilitates the success of chrCM.

**Figure 6.**
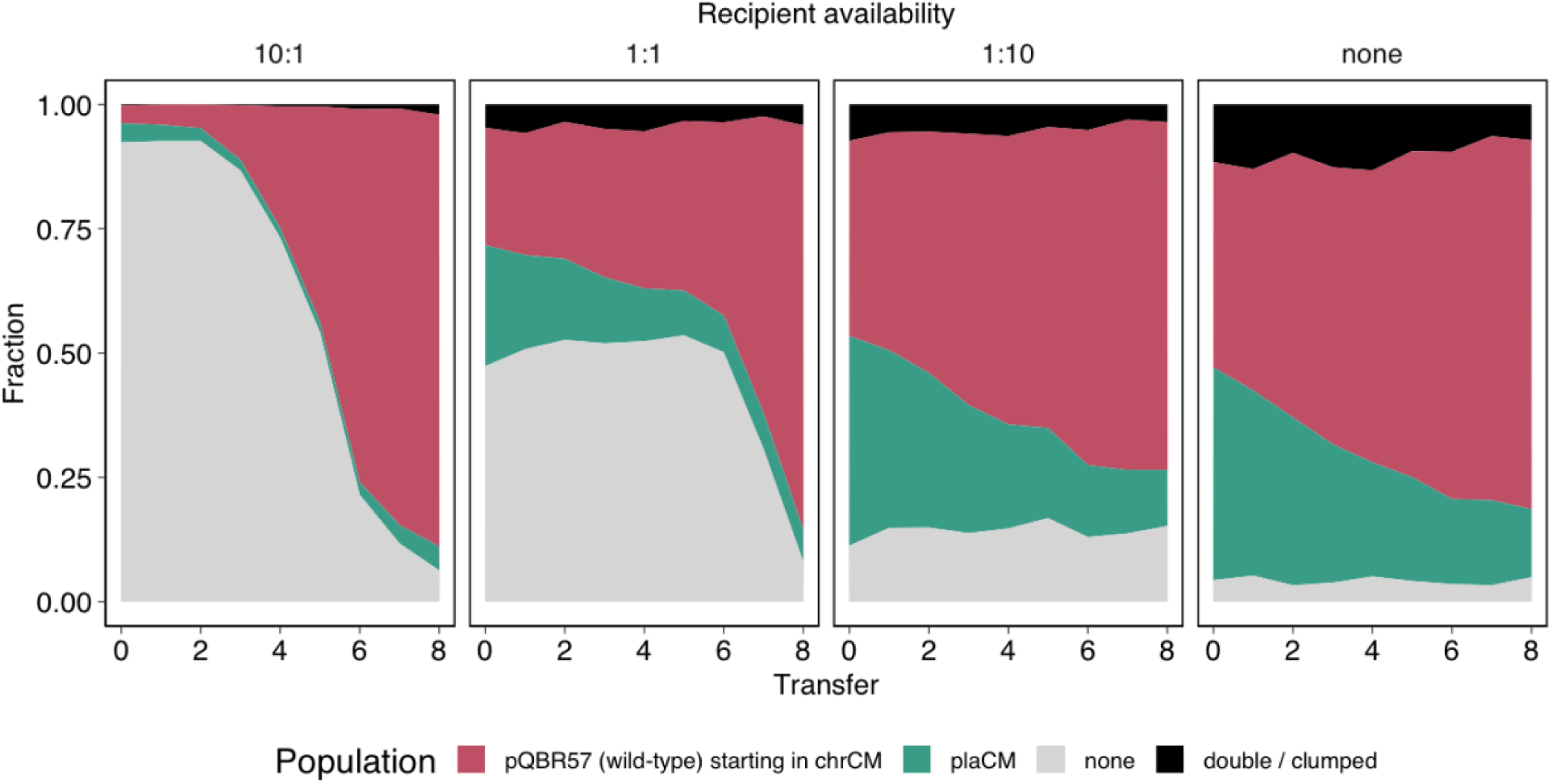
Plasmids carried by chrCM cells were more successful than plaCM plasmids regardless of recipient availability. Plots are arranged as Fig. 5. Plots indicate the mean of 6 independent experiments; replicate-level plots are provided in Figure S4.

The relative inability of plaCM to invade the recipient population, when compared with wild-type pQBR57 from chrCM cells, suggested that either plaCM exerted a negative pleiotropic effect on the ability of pQBR57 to invade a recipient population, or, conversely, that chrCM enhanced the ability of wild-type pQBR57 to invade. To distinguish between these possibilities, we conducted a similar experiment to Fig. 6, but here, plaCM was competed against pQBR57 harboured by an uncompensated competitor. As with Fig. 6, fluorescent labels were designed to track the relative success of the two plasmids. Unexpectedly, plaCM was only able to outcompete wild-type pQBR57 in the situation where plasmid-free recipients started at low frequency, i.e. effectively a head-to-head competition between compensated and uncompensated plasmid-bearers. We detected a significant effect of plasmid-free recipient ratio on the competition dynamics (GLMM fourth-order polynomial transfer:ratio interaction 𝜒^2^ = 414.4, p < 1e-7). Specifically, under conditions where recipients were more numerous, wild-type pQBR57 outcompeted the plaCM despite higher fitness costs (Fig. 2), an observation that suggests that plaCM inhibits plasmid transmission (or establishment in transconjugants) relative to the wild-type. Overall, plaCM did not appear to gain a substantive fitness benefit from horizontal transmission, only reaching dominance under conditions that prioritise vertical replication and only in populations without chrCMs.

Our analytical models for plaCM predicted cyclical RPS-like dynamics for some combinations of parameters (Fig 1B orange region). The experimental results in Fig. 7 were consistent with this prediction. Specifically, plasmid-free populations were invaded by the wild-type plasmid (left panel, and mid-left panel after transfer 4), the wild-type plasmid was outcompeted by plaCM (right panel, mid-right panel, and mid-left panel before transfer 4), and plaCM was outcompeted by plasmid-free (mid-right panel). To explore the dynamics in further detail we experimentally determined key parameters in our system (Table S1) and compared these with the analytical model (Fig. 1). With our system, and with experimentally-measured parameter values, the uncompensated plasmid exceeds the threshold for plasmid invasion and domination, which is indeed what we have described previously (Stevenson et al., 2017). Both chrCM and plaCM likewise exceed the threshold for domination by 18- and 10-fold respectively, indicating that, with sufficient reduction in 𝛾_𝑄_ relative to 𝛾_𝑃_, the plaCM system might exist in the (*f*\*,*p*\*,*q*\*) region of parameter space. Numerical simulations based on equations 1–6 and Supplementary Equations 1, and parameterised with biologically-plausible values, resulted in dynamics resembling observed experimental results, provided plaCM decreased conjugation rate sufficiently (10–100×, Fig. 8). An interactive (Shiny) app enabling readers to explore these patterns is provided at (<u>jpjh.shinyapps.io/COMPMOD_shiny</u>).

**Figure 7.**
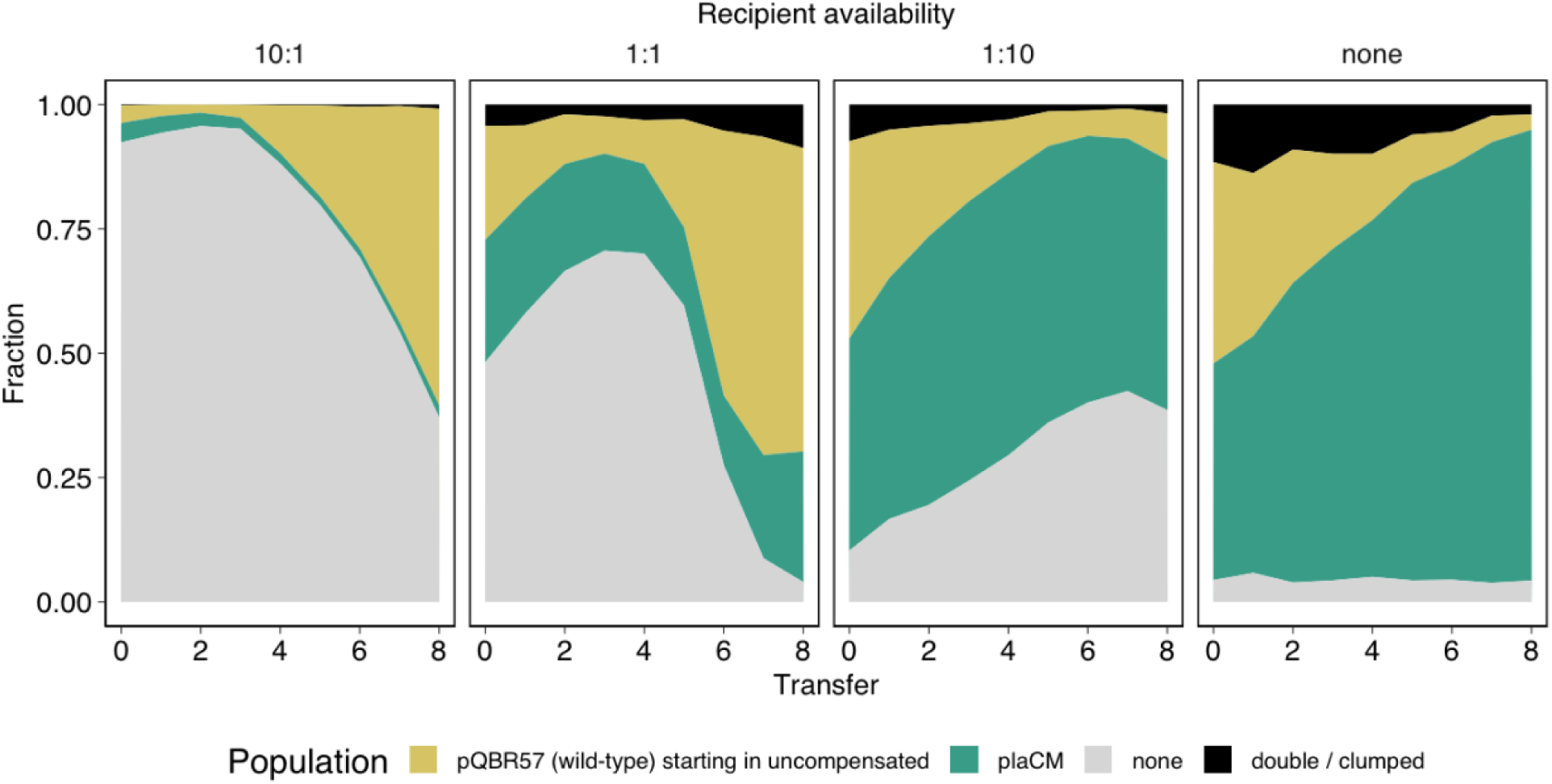
Plasmid-free, uncompensated, and plaCM-compensated plasmids undergo ‘rock-paper-scissors’ cyclical dynamics. Plots are arranged as Fig. 5. Plots indicate the mean of 6 independent experiments; replicate-level plots are provided in Figure S5.

**Figure 8.**
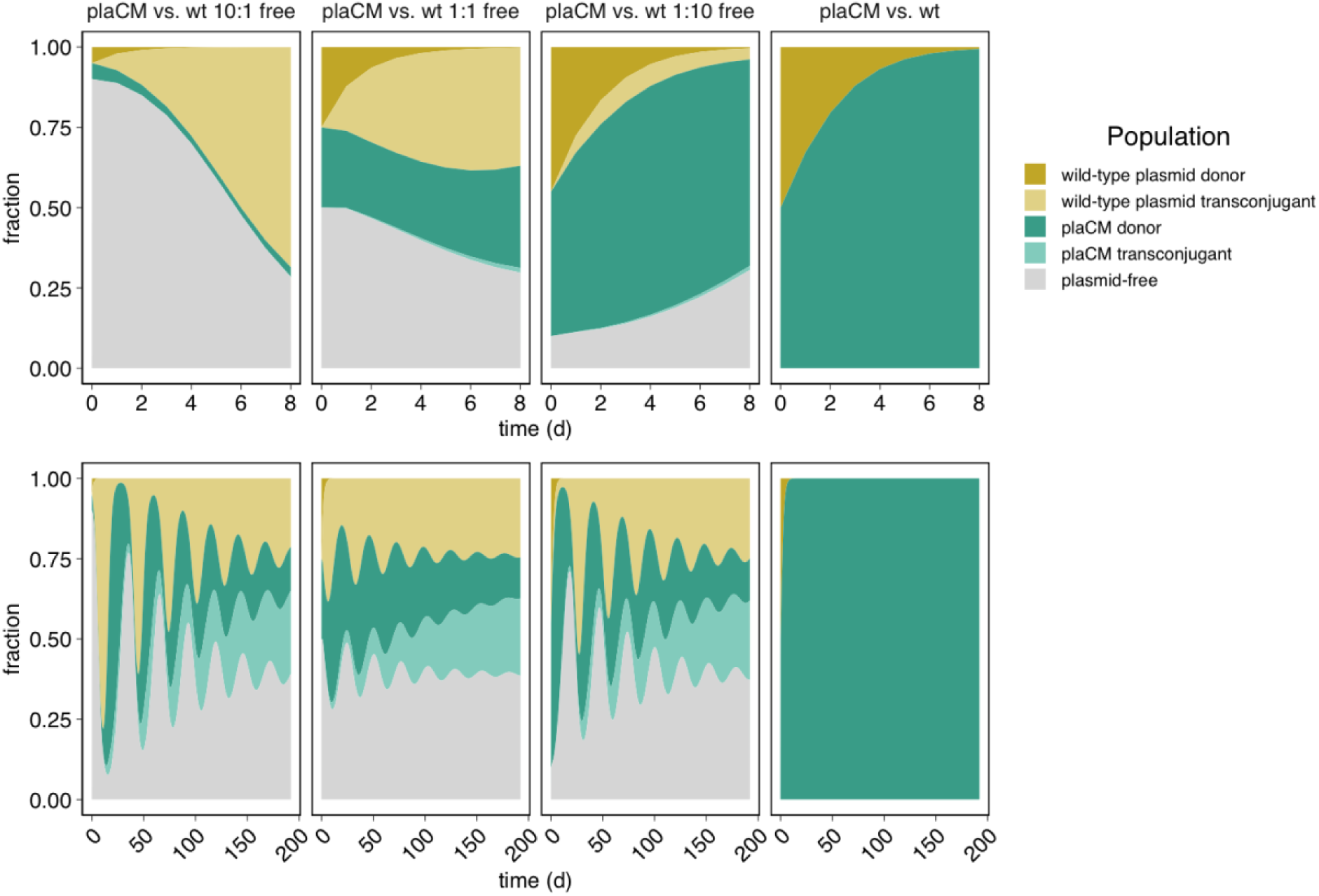
ODE-based model simulations resemble experimental results. Numerical simulations of a batch-transfer model using the following parameters: 𝛼 = 0.6 h^-1^, SBW25(pQBR57) relative fitness = 0.82, SBW25(pQBR57::plaCM) relative fitness = 0.95, *K* = 5.7 x 10^9^ ml^-1^, uncompensated conjugation rate = 9.9 x 10^-12^ ml.cells^-1^h^-1^, plaCM conjugation rate = 1.3 x 10^-13^ ml/cells/h; 𝛾_𝑃_ ∼ 75 × 𝛾_𝑄_. Top panels indicate dynamics over 8 transfers, bottom panels over 192 transfers. Details on parameterisation are provided in Supplementary Table 1. An interactive version of this figure is provided at <u>jpjh.shinyapps.io/COMPMOD_shiny</u>.

Together, our experimental and modelling results suggested that plaCM has a negative pleiotropic effect on conjugative transmissibility. Previous attempts to measure the intraspecific conjugation rates of pQBR57 with and without chrCM/plaCM did not detect any significant differences from the wild-type pQBR57 (Hall et al., 2021). However, these experiments were conducted over a relatively long time window (24 hr), which could allow fitness differences between transconjugants, donors, and recipients to mask differences in transfer rate (Huisman et al., 2022; Kosterlitz et al., 2022). We therefore re-measured conjugation rate using the Approximate Extended Simonsen approach (Huisman et al., 2022) — an extension to the popular ‘Simonsen’s gamma’ (Simonsen et al., 1990) that accommodates the potential for variation in growth rate. Unexpectedly, we did not detect any significant difference between the wild-type and plaCM variants of pQBR57, and certainly not the order-of-magnitude difference predicted by the model (Figure S6; t_9.86_ = 0.75, p = 0.94, TOST equivalence test with log_10_- transformed bounds ±0.5, p = 0.02). We therefore hypothesise that the differences in pQBR57 transmissibility between wild-type and plaCM variants observed in Fig. 7 emerge from yet-to-be-determined processes following plasmid establishment in recipient cells.

## Discussion

Plasmid fitness cost amelioration is an important process driving the maintenance, distribution, and dissemination of plasmids and their associated traits. Where plasmid fitness costs are generated from an interaction between the plasmid and resident chromosomal genes, mutations affecting either of these partners can enable plasmid survival. Previous experimental evolution studies on CMs have generally implicated chromosomal loci rather than plasmid loci as the principal targets, a finding which has been taken to reflect mutational supply, availability, and/or the poor ability of oftentimes recessive compensatory mutations to penetrate when appearing on a multi-copy plasmid (Mei et al., 2019; Stalder et al., 2017). But although more difficult to access, plaCMs, once achieved, ought to be more successful than chrCMs under a range of ecological conditions owing to the simple fact that plaCM is propagated when the plasmid transfers into recipients (Zwanzig et al., 2019). The relatively high transfer rate of pQBR57 ought to have further accentuated this benefit (Hall et al., 2015), particularly under environmental conditions with high recipient availability. In contrast to these expectations, our analyses and experiments showed that plaCM was not successful under most tested conditions, losing out to chrCM, wild-type plasmids harboured by chrCM-containing cells, and even, where opportunities for HGT were plentiful, wild-type plasmids from cells lacking CMs.

There are several processes that could explain the relative failure of plaCM. First, our analyses of extensions to a simple plasmid population dynamics model reveal that for transmissible plasmids, chrCMs provide a hidden benefit besides directly reducing the fitness cost of plasmid carriage to their bearers: conjugation from chrCM-containing cells transforms plasmid-free competitors into plasmid-carriers suffering the full burden of uncompensated plasmid carriage, indirectly enhancing the relative fitness of chrCM. Previous numerical simulations have likewise demonstrated the possibility for ‘weaponisation’ of conjugative elements through compensatory mutation, and such a dynamic provides a further explanation for the persistence of non-beneficial plasmids in communities (Domingues et al., 2022; Rebelo et al., 2023b, 2023a). Our analyses generalise these findings and demonstrate that a high degree of amelioration and low impact on transmissibility will enhance chrCM invasion, particularly if the local chrCM frequency is sufficiently high, a condition that is more likely to be met in spatially-structured habitats and/or where chrCM confers pleiotropic environmentally-adaptive benefits (Kloos et al., 2021; Loftie-Eaton et al., 2017; Rebelo et al., 2023a). Thus, the marginal benefits of conjugative transmissibility for plaCM in competition with chrCM are reduced.

Second, the benefits to plaCM of conjugative transmission wane as the plasmid-free recipient pool is diminished, from either (i) selection against plasmid-free recipients, or (ii) recipient acquisition of plasmids that can bar the incoming plaCM by incompatibility or exclusion. We observed both dynamics in our experiments. The effects of the latter mechanism are even more pronounced if there is a mechanistic trade-off between compensating plasmid fitness costs and the ability of the plasmid to transfer or establish in recipients, such as our results suggest, since a more transmissible (but more costly) plasmid can sweep through the recipient population, blocking access for the plaCM. Once the plasmid-free population is diminished, the dynamics of the system are driven by the relative growth rates of the different subpopulations, which in turn are determined by the degree of amelioration provided by chrCM and plaCM. In our system, SBW25::chrCM(pQBR57) outgrows SBW25(pQBR57::plaCM) in direct competition, because plaCM is not as efficient as chrCM at compensating the fitness cost of pQBR57. This discrepancy is likely due to the molecular mechanisms that underpin the principal fitness costs of pQBR57 and their resolution by compensatory mutation. PFLU4242 is a putative endonuclease, with a DUF262/DUF1524 domain structure resembling that of the GmrSD type IV restriction system (Machnicka et al., 2015), and so we hypothesised that this gene is somehow directly responsible for generating the dsDNA breaks that trigger the SOS response and subsequent toxic gene expression patterns characteristic of uncompensated pQBR57 carriage (Hall et al., 2021). Loss-of-function mutations to PFLU4242 (i.e. chrCM) would directly prevent these breaks from occurring. On the other hand, *PQBR57_0059* encodes a lambda repressor-like protein that regulates expression of two other pQBR57-encoded putative DNA-binding proteins, PQBR57_0054-0055, and it is upregulation of PQBR57_0054-0055 which provides the proximal mechanism of plaCM compensation, through a mechanism not yet fully understood. ChrCM therefore likely provides a more direct route than plaCM to resolving the genetic conflict at the heart of pQBR57-SBW25 fitness costs, and thus is mechanistically a more effective CM. Ultimately, it is likely to be the degree to which CMs reduce the cost of plasmid carriage, rather than CM transmissibility, which will primarily determine CM success.

Unexpectedly, plaCM was a poor competitor against the wild-type uncompensated pQBR57, winning out only in cases where there were few opportunities for conjugation. This experimental observation, coupled with our model parameterisation and numerical simulations, strongly implies that pQBR57::plaCM is not as effective as the wild-type plasmid at transmitting by conjugation and/or establishing in recipients. As measured conjugation rates of wild-type pQBR57 and pQBR57::plaCM have been indistinguishable, even when controlling for the different fitness effects of the two plasmids, these observations imply that differences in transmissibility emerge from processes other than plasmid transfer *per se*. PQBR57_0055 is a Spo0J/ParB-like protein, homologues of which have been shown to have various, non-specific effects on gene expression, and by upregulating PQBR57_0054-0055, plaCM could have various pleiotropic effects in transconjugants. For example, plasmids have been shown to impose transient costs on acquisition, mainly driven by increased in lag time and usually resolved in a matter of hours, alongside the longer lasting fitness costs (Ahmad et al., 2023; Prensky et al., 2021). Additionally, some plasmids exhibit ‘conjugation derepression’, whereby, for a short period, transconjugants display an increased onward conjugation rate. The extent and duration of such ‘acquisition costs’ and conjugation derepression may vary between plaCM and wild-type to disfavour plaCM transmission without having a direct impact on transmission rate. Overall, our experiments comparing pQBR57::plaCM and wild-type pQBR57 are congruent with several other studies demonstrating trade-offs between vertical and horizontal plasmid transmission (Bethke et al., 2023; Dahlberg and Chao, 2003; Dimitriu et al., 2021; Porse et al., 2016; Turner et al., 2014), and further show that imperfect plaCMs that trade-off against transmission can generate long-standing oscillatory dynamics which could sustain diversity in the plasmid population.

The chrCM in our system ameliorates diverse other mercury resistance plasmids, including pQBR103 and pQBR55 (Hall et al., 2021, 2019), and can ameliorate the costs of co-habiting compatible plasmids (Carrilero et al., 2020). Previous work has likewise shown the generality of chromosomal compensatory mutations in reducing the fitness costs of different plasmids (Loftie-Eaton et al., 2017). Our experiments did not detect a beneficial effect on chrCM vs. plaCM when pQBR103 was introduced, likely because the low conjugation rate of pQBR103, overall benefit of chrCM, and short period of the experiment meant that any selective pressure imposed by pQBR103 acquisition was negligible. Nevertheless, the fact that chrCM was more likely to outcompete plaCM even in a single-plasmid (pQBR57) system indicates that lineages gaining CMs can become pre-disposed to acquiring further plasmids, potentially becoming hubs for horizontal gene transfer, plasmid recombination, and trait dissemination in microbial communities. Efforts to limit HGT, for example to control the spread of antibiotic resistance, should therefore focus on identifying and targeting such ‘keystone’ strains. One possible route would be to use antagonistic parasitic MGEs such as lytic bacteriophage. The target of chrCM in our system appears related to GmrSD, a known genome defence mechanism, and while little was known of the biological function of the *P. aeruginosa* PAO1 chrCM targets identified by San Millan et al. (San Millan et al., 2015) at the time of discovery, gene function prediction tools now associate the accessory helicase PA1372 and partner gene PA1371 with genome defence (‘Helicase + DUF2290 system’) (Payne et al., 2021; Tesson et al., 2022). Likewise, the ‘Xpd/Rad3-like helicase’ and ‘upstream UvrD helicase’ targets of chrCMs identified by Loftie-Eaton et al. (Loftie-Eaton et al., 2017) in *Pseudomonas* sp. H12 refer to predicted components of prokaryotic Argonaute type III and Gabija respectively, while in *Vibrio*, a recently-discovered defence system DdmABC confers a high fitness cost on bearers of large plasmids such that they are removed from a population by purifying selection in a manner that resembles the large fitness cost imposed by PFLU4242 (Jaskólska et al., 2022). The ability of ‘MGE-favourable’ organisms to receive and host plasmids thus likely trades off against susceptibility to costly parasitic elements, and exploiting this weakness may be a profitable approach to controlling the maintenance and spread of unwanted mobile genetic elements in various microbiomes.

Though some aspects — namely the relative degree to which chrCM and plaCM ameliorate plasmids and the extent to which plaCM imposes pleiotropic effects on conjugation rate — may be system specific, the superiority of chromosomal CMs over plasmid-borne CMs in terms of mutational accessibility, indirect effects on plasmid-free competitors (as presented here with a general theoretical mechanism), mechanistic efficacy, and reduced potential for trade-off with horizontal transmission, are likely to generalise to diverse other plasmid-bacterial pairings, including pathogens and multi-drug resistance plasmids. One broader implication is that transmissible plasmids thus have a limited ability to ‘act nice’ by effectively ameliorating their own costs by evolution. Instead, plasmids are under stronger selection to improve their transmissibility and intracellular competitiveness, and it is largely down to resident genes to accommodate these unruly vectors — or remove them.

## Acknowledgements

This work was supported by the Natural Environment Research Council (NE/R008825/1) to MAB, EH, AJW, SP, JPJH. RCW is supported by a funding from the Biotechnology and Biosciences Research Council to RCW and MAB (BB/T014342/1). JPJH is supported by an MRC Career Development Award (MR/W02666X/1). MJB is supported by the Wellcome Trust Sir Henry Wellcome Fellowship (221663/Z/20/Z).

## Author contributions

Conceptualization: RCTW, AJW, SP, EH, MAB, JPJH. Data curation: RCTW, JPJH, Investigation: RCTW, AJW, MJB, JPJH. Resources: RCTW, KJM, JPJH. Formal analysis: RCTW, AJW, MJB, JPJH. Funding acquisition: MAB, EH, AJW, SP, JPJH. Visualization: RCTW, AJW, MJB, JPJH. Writing — original draft: RCTW, AJW, MAB, JPJH. Writing — review & editing: RCTW, AJW, MJB, EH, MAB, JPJH.

## Methods

### Bacterial strains

Fluorescently-labelled strains of *Pseudomonas fluorescens* SBW25 and *P. fluorescens* SBW25ΔPFLU4242 (Hall et al., 2019) were generated using the mini-Tn7 system and plasmid pUCT-mini-Tn7T-Gm-eyfp (Choi and Schweizer, 2006) or a derivative in which dTomato was cloned to replace eyfp. The pQBR57ΔPQBR57_0059 knockout, and fluorescently-labelled variants of megaplasmid pQBR57 (Hall et al., 2015; Lilley et al., 1996) were generated using homologous recombination with plasmid pTS-1 (Campilongo et al., 2017). Briefly, for the knockout, 1 kb flanking regions of pQBR57 were amplified and cloned into the MCS of Xba-KpnI-digested pTS-1 using NEB HiFi assembly. For the fluorescently-labelled plasmids, fragments of pQBR57 and an expression cassette consisting of a Ptac promoter driving a fluorescence protein gene (either eGFP or tdTomato), followed by lambda t0 and rrnB T1 terminators were amplified and cloned into the MCS of XhoI/KpnI-digested pTS-1 using NEB HiFi assembly. Constructs were transformed into SBW25(pQBR57) by electroporation (Choi and Schweizer, 2006), and merodiploids selected on KB supplemented with 100 µg/ml tetracycline. Double-crossovers were selected on LB supplemented with 10% w/v sucrose and 20 µM HgCl_2_, and candidates screened by PCR and tetracycline sensitivity before sending for whole genome sequencing (2x250 bp, >30x coverage, MicrobesNG) to test for second-site mutations. Using breseq (Deatherage and Barrick, 2014), no second-site mutations were detected for the strains used in these experiments. For experiments, each plasmid-containing replicate was established with an independent transconjugant from a genome-sequenced donor strain. Conjugation experiments were performed with non-fluorescent antibiotically-labelled strains described previously (Hall et al., 2021).

### Competition experiment

To determine the short-term fitness effects of compensation, direct competitions were performed between YFP- *versus* dTomato-labelled strains. Overnight cultures were mixed at 1:1 ratio (test:reference) before inoculation at 1:100 dilution into 6 ml King’s B media in a 30 ml glass universal with loose-fitting lid (‘microcosm’), with or without mercury (Hg (II), 40 µM) and incubated at 28°C, 180 rpm for 24 hours. Each competition was repeated with 10 biological replicates. Flow cytometry was used to estimate bacterial counts: starting mixtures and endpoint competition cultures were diluted 1:100 into M9 buffer and run on a Beckman Coulter CytoflexS machine at 17 µl.min^-1^ for either 5,000 counts (determined as events with signal in SSC-H channel > 10^3^) or 90 seconds maximum. Between samples, M9 buffer was sampled for 5 seconds to minimise cross-over between samples. Strain counts were determined by gating in the following channels: FITC-H (gate=10^3.4^ for YFP) and PE-H (gate=10^3.4^ for dTomato). Relative fitness was calculated as the difference in Malthusian parameters, 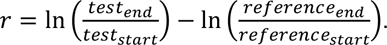 .

### Serial passage experiments

For all evolution experiments, bacterial populations were grown in 6 ml KB microcosms at 28 °C with agitation at 180 rpm. Serial daily transfers of 1% population into fresh media were performed for 8 days, with daily flow cytometry used to track population dynamics. For flow cytometry, cultures were diluted 1:100 into M9 buffer and incubated with Hoechst 34580 stain (5 µg/ml) for 15 minutes in the dark at room temperature to enable detection of unlabelled bacterial strains. Flow cytometry data was sampled for 60 seconds at 17µl.min^-1^ with minimal gating for size (FSC-H > 10^3^), and strain counts were determined in post analysis with the following thresholds: Hoechst-stained bacteria (i.e., total bacterial count, PB450-H > 10^4^), YFP only (FITC-H > 10^3.5^), dTomato only (PE-H > 10^3.4^), YFP+dTomato clumps (FITC-H > 10^3.5^ and PE-H > 10^3.4^). Between samples, M9 buffer was sampled for 5 seconds to minimise cross-over between samples. For each fluorescently labelled strain, single-strain populations (3 biological replicates) were serially transferred for the duration of the experiment to ensure maintenance of the fluorescent signal.

To investigate the benefit of differing modes of compensation under varying selection pressures, we first challenged equal proportions of plaCM (SBW25(pQBR57Δ0059)) against chrCM (SBW25ΔPFLU4242(pQBR57)) in the presence of either a plasmid-free wild-type host (SBW25) or a wild-type host bearing a more costly conjugative plasmid (SBW25(pQBR103)). In all populations, the wild-type host started at ∼50% frequency and carried a gentamicin resistance marker (Gm^R^). In a fully factorial design, each population was grown in either in the presence or absence of mercury (Hg (II), 40 µM). Fluorescent markers (YFP or dTomato) associated with each mode of compensation allowed tracking of population dynamics in the presence of a non-fluorescent wild-type strain, with 10 biological replicates per marker orientation.

The impact of host availability on the benefit of plaCM was investigated in a separate evolution experiment, by challenging GFP-labelled plaCM (SBW25(pQBR57Δ0059::GFP)) against different host:plasmid backgrounds in the presence of varying ratios of plasmid-free wild-type hosts (SBW25::Gm^R^), including 10x excess, equal proportions, 10x fewer and no available hosts. At each level of host availability, plaCM was competed against chrCM with a chromosomally-encoded fluorescent label (SBW25Δ4242::dTomato(pQBR57)), chrCM with a plasmid-encoded fluorescent label (SBW25Δ4242(pQBR57::tdTomato)) or a wild-type plasmid bearer (SBW25(pQBR57::tdTomato)). For each treatment, 6 biological replicates were performed. Raw data from flow cytometry experiments are provided at https://doi.org/10.5285/51046841-deaa-422f-a303-2c0759f014b4.

### Conjugation rates

Streptomycin-resistant *lacZ*-carrying donors (either SBW25(pQBR57) or SBW25(pQBR57ΔpQBR57_0059)), Gm^R^ recipients (SBW25), and Gm^R^ transconjugants (either SBW25(pQBR57) or SBW25(pQBR57ΔpQBR57_0059)), cultured overnight in 150 µl KB broth in an untreated CytoOne 96-well microtitre plate at 28°C, were subcultured 1:30 into 150 µl fresh media and placed in a Tecan Nano plate reader for incubation at 28°C with shaking. When exponential phase was reached (assessed by examination of growth curves; OD600 ∼ 0.4) cultures were again diluted 30-fold into KB. Mixed cultures containing donors and recipients, or single-strain donor, recipient, or transconjugant cultures were sampled and cultured in the plate reader and spread on KB agar plates supplemented with 50 µg/ml X-gal for enumeration. After approx. 4 hours growth, cultures were again sampled and spread on KB agar plates, some of which were supplemented with antibiotics (250 µg/ml streptomycin or 30 µg/ml gentamicin) and 20 µM mercury to enumerate transconjugants. Conjugation rates were calculated with the Approximate Extended Simonsen Method (Huisman et al., 2022).

### Statistics

Relative fitness was analysed using linear models, with post-hoc pairwise comparisons performed using the package emmeans (Lenth, 2023). Dynamics were analysed with Generalized Linear Mixed Effects Models (GLMM) using the R package glmmTMB (Brooks et al., 2017), with a beta-binomial response distribution, a logit link function, and the counts of each competitor as the response variables. Preliminary analyses identified overdispersion in the data, justifying the use of a beta-binomial rather than a binomial response distribution (Harrison, 2015). Non-independence of measurements arising from repeated sampling of populations was accommodated with random effects of ‘population’ on intercept and slope, except for the experiment presented in Fig. 7 which included only the random effect on intercept due to extremely low variance and high correlation between random effects preventing model convergence. For experiments presented in Figs. 6 and 7, polynomial terms were added to accommodate potentially non-monotonic relationships between competitors over time. Significance of fixed effects were determined by comparison of nested models using likelihood ratio tests. Conjugation rate data were analysed by t-test and two one-sided tests using the package TOSTER (Lakens et al., 2018). Data and analysis scripts are provided at https://github.com/jpjh/COMPMUT_dynamics.

