## Supplementary Information for "Superiority of chromosomal compared to plasmid-encoded compensatory mutations"

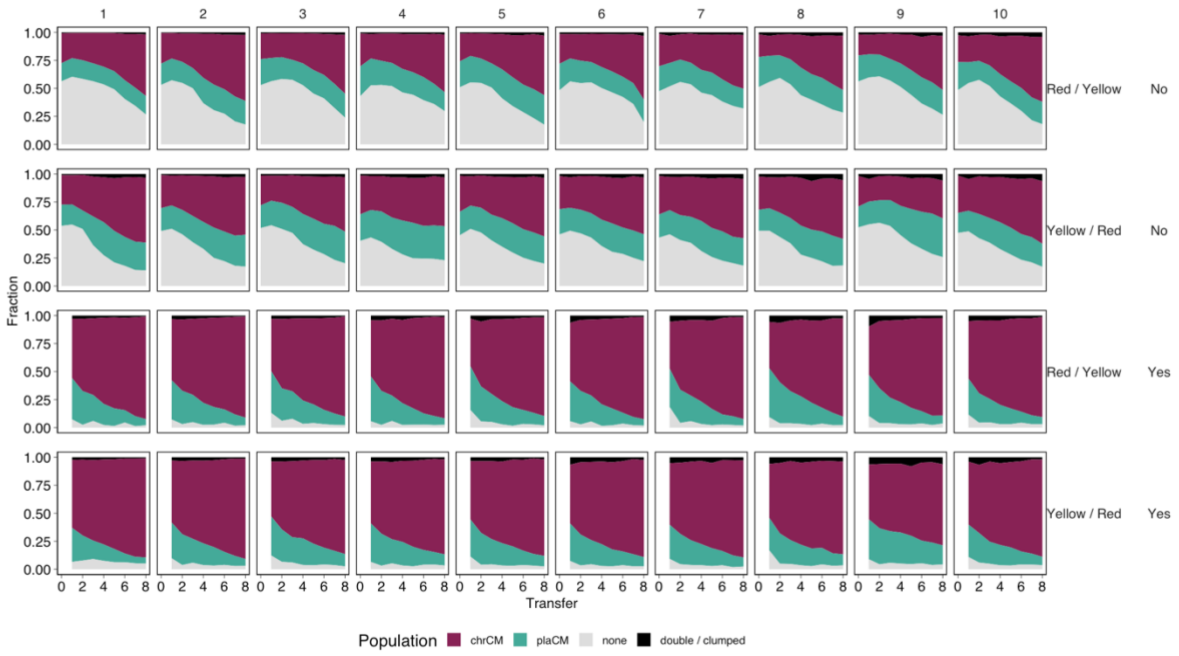


**Figure S1.** Individual replicates for the summarized data presented in Figure 3. Red/Yellow and Yellow/Red refer to the orientation of the fluorescent markers (chrCM / plaCM), and Yes / No refers to mercury selection.

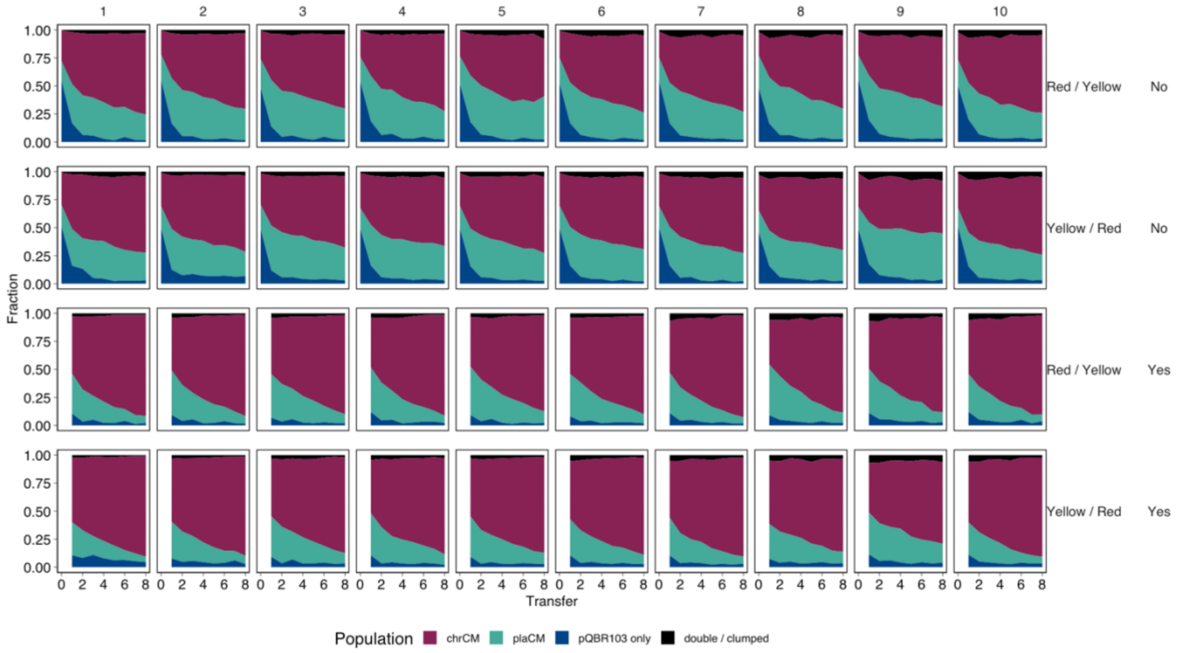


**Figure S2.** Individual replicates for the summarized data presented in Figure 4. Red/Yellow and Yellow/Red refer to the orientation of the fluorescent markers (chrCM / plaCM), and Yes / No refers to mercury selection.

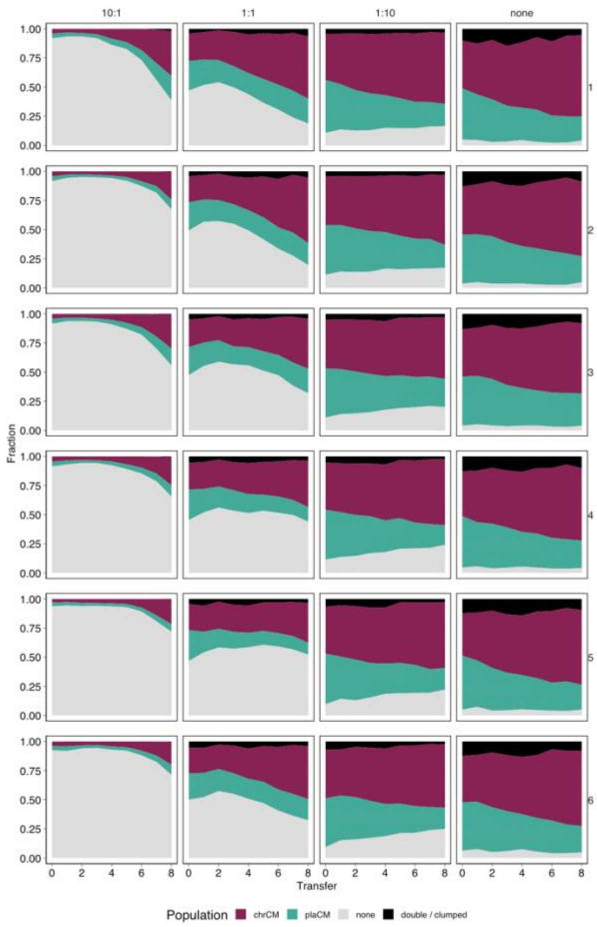


**Figure S3.** Individual replicates for the summarised data presented in Figure 5.

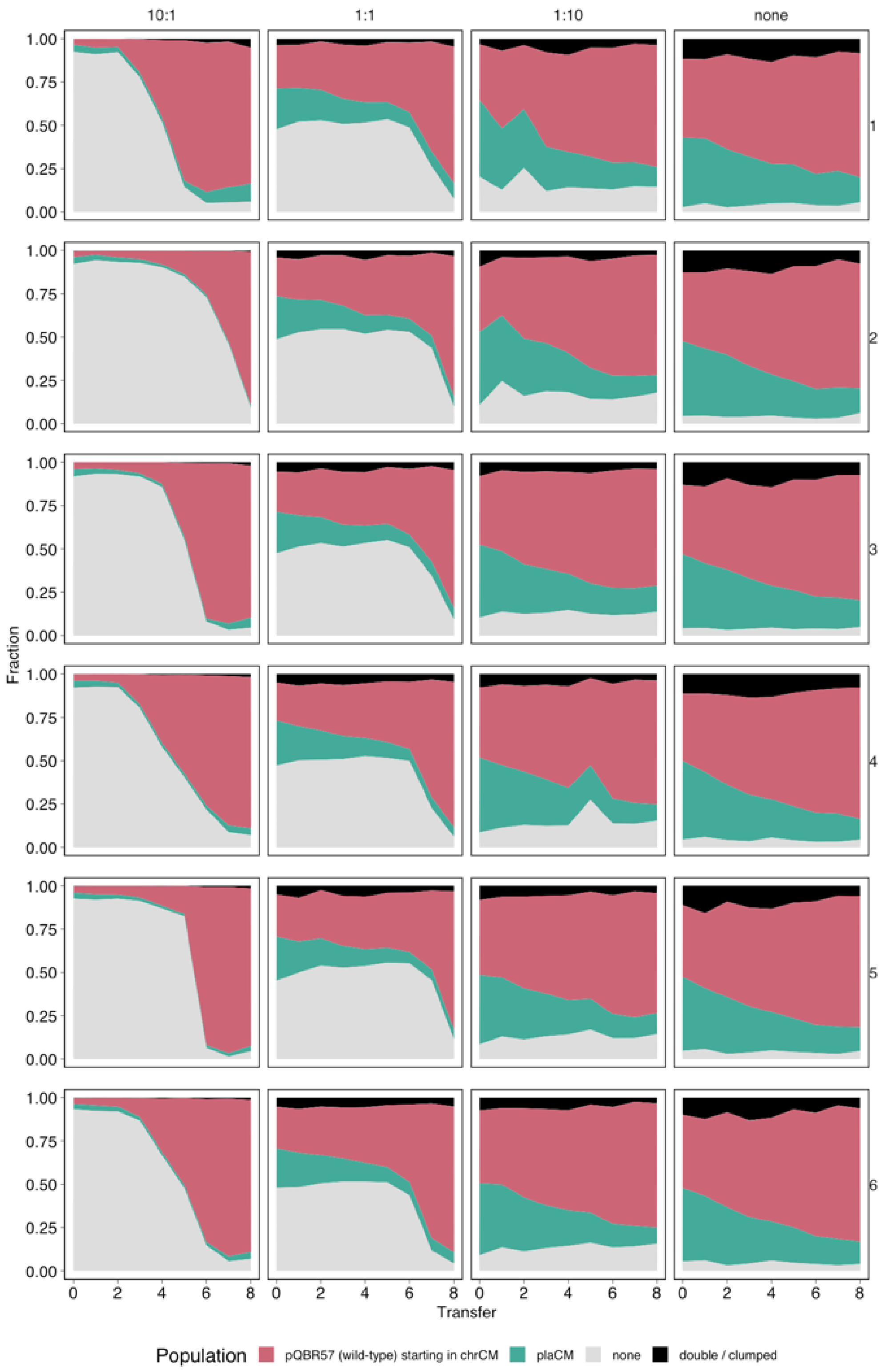


**Figure S4.** Individual replicates for the summarised data presented in Figure 6.

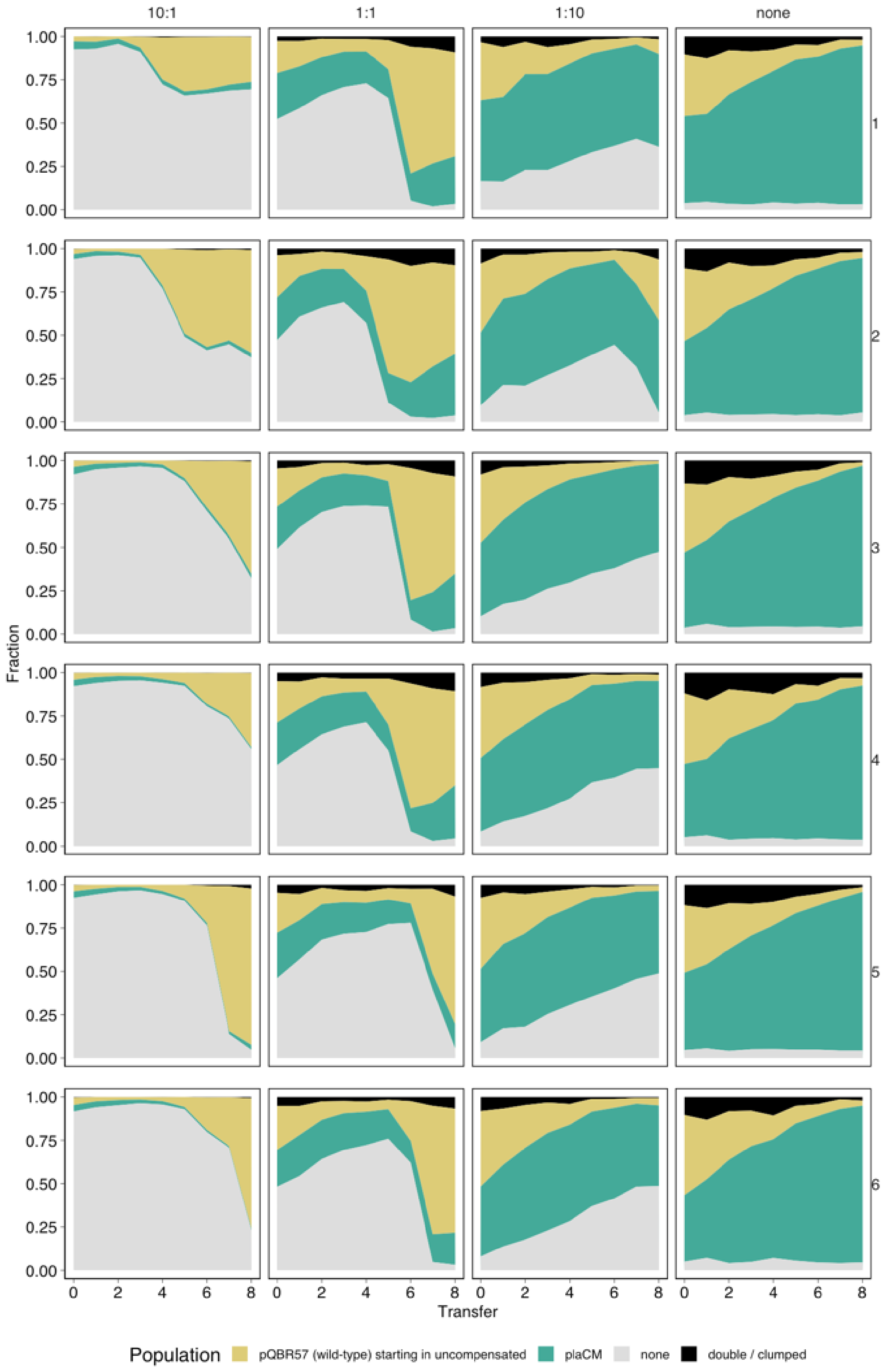


**Figure S5.** Individual replicates for the summarised data presented in Figure 7.

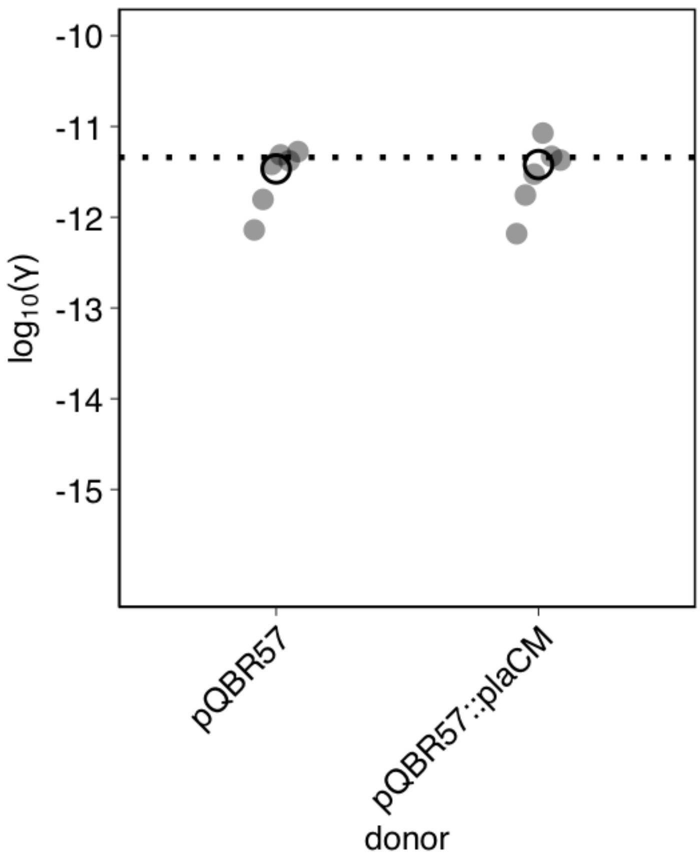


**Figure S6.** Approximate Extended Simonsen conjugation rates for pQBR57 and pQBR57::plaCM. The dotted line indicates the previously-measured conjugation rate for wild-type pQBR57 (Hall et al. 2021, Table S1).

**Table S1. Experimentally measured parameters used to inform numerical simulations**

| Parameter* | Units | Value | Standard Deviation | Notes and source data |
|---|---|---|---|---|
| $\alpha$ | $h^{-1}$ | 0.541 | 0.0553 | Calculated using gcplyr (Blazanin 2023) from growth curve data collected by subculturing overnight cultures 1:1000 into 150 µl fresh KB media and measuring optical density at 600 nm using a Tecan Nano plate reader every 15 minutes for 24 hours. Mean of 12 replicate cultures. |
| SBW25(pQBR57) relative fitness | dimensionless | 0.818 | 0.0429 | (Hall et al., 2021). Multiply by $\alpha$ for $\beta_P$. |
| SBW25(pQBR57::plaCM) relative fitness | dimensionless | 0.969 | 0.0699 | (Hall et al., 2021). Multiply by $\alpha$ for $\beta_Q$. |
| SBW25::chrCM(pQBR57) relative fitness | dimensionless | 0.945 | 0.0413 | (Hall et al., 2021). Multiply by $\alpha$ for $\beta_C$. |
| K | $ml^{-1}$ | 5.72e9 | 1.59e9 | (Hall et al., 2021) |
| pQBR57 conjugation rate | ml/cells/h | 4.57e-12 | 1.014e-12 | (Hall et al., 2021). Multiply by $K$ to give $\gamma$ in the (scaled) analytic model. |
| $\mu$ | $h^{-1}$ | 0.04125 | 0 | Calculated as an average considering a 1:100 dilution/24h. |

**Supplementary Text 1**

The following equations describe the numerical simulation model presented at jpjh.shinyapps.io/COMPMOD_shiny/ and in Figure 8. This model has been generalised to flexibly describe different types of compensatory mutation (CM). Here, $Z_f$ describes the wild-type plasmid-free population, $X_0$ describes the wild-type population with the wild-type plasmid, $X_1$ describes a plasmid-containing population with CM type 1, and $X_2$ describes a plasmid-containing population with CM type 2. $X_{0t}$, $X_{1t}$, and $X_{2t}$ describe transconjugants formed by transfer into $Z_f$ from $X_0$, $X_1$, and $X_2$ respectively. Each population has its own growth rate, conjugation rate, and susceptibility to positive selection, and by adjusting these parameters, the features of plaCM and chrCM can be described. Specifically, to describe a plaCM, the growth rate of the transconjugant is set to be the same as that of the donor (e.g. $\alpha_{X_{1t}} = \alpha_{X_1}$), whereas to describe a chrCM, the growth rate of the transconjugant is set to be the same as that of $X_0$, the wild-type population with the wild-type plasmid (e.g. $\alpha_{X_{2t}} = \alpha_{X_0}$).

$$\sigma = 1 - \frac{(Z_f + X_0 + X_1 + X_2 + X_{0t} + X_{1t} + X_{2t})}{K}$$

$$\frac{dZ_f}{dt} = \alpha_{Z_f}\sigma Z_f - \mu Z_f - \gamma_{X_0}X_0Z_f - \gamma_{X_1}X_1Z_f - \gamma_{X_2}X_2Z_f - \gamma_{X_{0t}}X_{0t}Z_f - \gamma_{X_{1t}}X_{1t}Z_f - \gamma_{X_{2t}}X_{2t}Z_f - \eta_{Z_f}Z_f$$

$$\frac{dX_0}{dt} = \alpha_{X_0}\sigma X_0 - \mu X_0 - \eta_{X_0}X_0$$

$$\frac{dX_1}{dt} = \alpha_{X_1}\sigma X_1 - \mu X_1 - \eta_{X_1}X_1$$

$$\frac{dX_2}{dt} = \alpha_{X_2}\sigma X_2 - \mu X_2 - \eta_{X_2}X_2$$

$$\frac{dX_{0t}}{dt} = \alpha_{X_{0t}}\sigma X_{0t} - \mu X_{0t} + \gamma_{X_0}X_0Z_f + \gamma_{X_{0t}}X_{0t}Z_f - \eta_{X_{0t}}X_{0t}$$

$$\frac{dX_{1t}}{dt} = \alpha_{X_{1t}}\sigma X_{1t} - \mu X_{1t} + \gamma_{X_1}X_1Z_f + \gamma_{X_{1t}}X_{1t}Z_f - \eta_{X_{1t}}X_{1t}$$

$$\frac{dX_{2t}}{dt} = \alpha_{X_{2t}}\sigma X_{2t} - \mu X_{2t} + \gamma_{X_2}X_2Z_f + \gamma_{X_{2t}}X_{2t}Z_f - \eta_{X_{2t}}X_{2t}$$

*(1)*

#### Supplementary Text 2 (Appendix)

### 1 Basic Model

The starting point is to first analyse the well known pair of dimensionless equations

$$\begin{aligned}\frac{df}{dt} =& \alpha f(1-f-p) - \mu f - \gamma p f \\ \frac{dp}{dt} =& \beta p(1-f-p) - \mu p + \gamma p f,\end{aligned} \tag{1}$$

which represent the growth of a plasmid free strain, $f$, and a plasmid containing strain, $p$ in the absence of selection. The strains have grown rate $\alpha$ and $\beta$. There is a washout rate $\mu$ and a conjugation rate $\gamma$. We take $\alpha > \beta > \mu$. A Jacobian matrix an be calculated to determine stability and is

$$J(f,p) = \begin{bmatrix} \alpha(1-2f-p) - \mu - \gamma p & -\alpha f - \gamma f \\ -\beta p + \gamma f & \beta(1-f-2p) - \mu + \gamma f \end{bmatrix} \tag{2}$$

This system has 4 fixed points which are

- $(0,0)$, this fixed point – no bacteria – is always unstable due to our assumptions
- $(f^*,0)$, this fixed point with $f^* = 1 - \dfrac{\mu}{\alpha}$ – no plasmid – is stable when $\gamma < \frac{\mu(\alpha-\beta)}{\alpha-\mu}$
- $(0,p^*)$, this fixed point with $p^* = 1 - \dfrac{\mu}{\beta}$– plasmid dominates – is stable when $\gamma > \frac{\mu(\alpha-\beta)}{\beta-\mu}$
- $(f^*,p^*)$, this fixed point with $p^* = \frac{\alpha}{\gamma z} - \frac{\mu}{\gamma}$ and $f^* = \frac{\mu}{\gamma} - \frac{\beta}{\gamma z}$ (where we define $z = \alpha - \beta + \gamma$) – a mixed solution – is stable when $\frac{\mu(\alpha-\beta)}{\alpha-\mu} < \gamma < \frac{\mu(\alpha-\beta)}{\beta-\mu}$

This gives a clear explanation for the expected dynamics as $\gamma$ varies: for $\gamma$ small conjugation is insufficient for the plasmid containing strain to invade the faster growing plasmid free strain and is eliminated. For large $\gamma$ the opposite occurs and the plasmid containing strain is able to infectiously invade and dominate the population, eliminating the plasmid free strain. For a range of intermediate values of $\gamma$ given by the inequality above, a balance is achieved between the growth rate advantage of the plasmid frees and them achieving sufficient levels in the population that they become infected with plasmids leading to coexistence.

### 2 Plasmid compensation model

The first variant we consider is if another strain emerges which is assumed to carry a mutation that compensates the cost of the plasmid somewhat. The new strain has a population fraction $q$ and its properties are subscripted $Q$, as apposed to $P$ for the original strain. The new plasmid has a differing conjugation rate $\gamma_Q$ as compared to $\gamma_P$. The full system has dynamics

$$\begin{aligned}\frac{df}{dt} =& \alpha f(1-f-p-q) - \mu f - \gamma_P pf - \gamma_Q qf \\ \frac{dp}{dt} =& \beta_P p(1-f-p-q) - \mu p + \gamma_P pf \\ \frac{dq}{dt} =& \beta_Q q(1-f-p-q) - \mu q + \gamma_Q qf, \end{aligned} \tag{3}$$

To match our assumptions we have chosen $\beta_P < \beta_Q \leq \alpha$ (WLOG mathematically). If the new conjugation rate $\gamma_Q \geq \gamma_P$ then the new strain is simply better than the old and will quickly displace it and we return to the basic model first described (with $p$ exchanged with $q$). Therefore interesting dynamics are only possible for $\gamma_Q < \gamma_P$. Taking these inequalities forward we find the following Jacobian matrix and fixed points:

$$J(f,p,q) = \begin{bmatrix} \alpha(\Delta - f) - \mu - \gamma_P f - \gamma_Q q & -\alpha f - \gamma_P f & -\alpha f - \gamma_Q f \\ -\beta_P p + \gamma_P p & \beta_P(\Delta - p) - \mu + \gamma_P f & -\beta_P p \\ -\beta_Q c + \gamma_Q q & -\beta_Q q & \beta_Q(\Delta - q) - \mu + \gamma_Q f \end{bmatrix} \tag{4}$$

with $\Delta = 1 - f - p - q$.

- $(0,0,0)$, this fixed point – no bacteria – is always unstable due to our assumptions
- $(f^*,0,0)$, this fixed point, $f^* = 1 - \dfrac{\mu}{\alpha}$ – no plasmid – is stable when $\gamma_P < \frac{\mu(\alpha-\beta_P)}{\alpha-\mu}$ and $\gamma_Q < \frac{\mu(\alpha-\beta_Q)}{\alpha-\mu}$
- $(0,p^*,0)$, this fixed point, $p^* = 1 - \dfrac{\mu}{\beta_P}$ – original plasmid dominates – is never stable because of our assumption that $\beta_Q > \beta_P$.
- $(0,0,q^*)$, this fixed point, $q^* = 1 - \dfrac{\mu}{\beta_Q}$ – new plasmid dominates – is stable when $\gamma_Q > \frac{\mu(\alpha-\beta_Q)}{\beta_Q-\mu}$
- $(0,p^*,q^*)$, this fixed point – a mixed plasmid solution – never exists due to our assumption that $\beta_P \neq \beta_Q$

In addition there are three further fixed points for which analytic progress is more challenging

- $(f^*,p^*,0)$, a fixed point – a mixed solution between frees and original plasmids.

- $(f^*, 0, q^*)$, a fixed point – a mixed solution between frees and new plasmids.
- $(f^*, p^*, q^*)$, a fixed point – a fully mixed solution.

The first two fixed points are mathematically symmetric to each other. In all cases interchanging $P \leftrightarrow Q$ will give the conditions for $(f^*, p^*, 0)$. We consider $(f^*, 0, q^*)$ WLOG, where $f^* = \frac{\mu}{\gamma_Q} - \frac{\beta_Q}{\gamma_Q z_Q}$ and $q^* = \frac{\alpha}{\gamma_Q z_Q} - \frac{\mu}{\gamma_Q}$ and we define $z_Q = \frac{\alpha - \beta_Q + \gamma_Q}{\gamma_Q}$ for convenience. This fixed point exists in the band $\frac{\mu(\alpha-\beta_Q)}{\alpha-\mu} < \gamma_Q < \frac{\mu(\alpha-\beta_Q)}{\beta_Q-\mu}$. Computing the eigenvalues of the Jacobian matrix leads to factorisation of the characteristic polynomial to the eigenvalue

$$\lambda_Q = \mu\frac{(\gamma_P - \gamma_Q)}{\gamma_Q} - \frac{\Omega}{\gamma_Q z_Q} \quad \text{where} \quad \Omega = \beta_Q\gamma_P - \beta_P\gamma_Q \tag{5}$$

and a quadratic. Solving the quadratic is not revealing but the Routh-Horwitz conditions lead to conditions corresponding to the existence conditions. Therefore this fixed point is stable when it exists and when $\lambda_Q < 0$. The fully mixed fixed point $(f^*, p^*, q^*)$, where $f^* = \frac{\mu(\beta_Q-\beta_P)}{\Omega}$, $p^* = \frac{\mu\gamma_Q z_Q}{\Omega} - \frac{\gamma_Q}{\gamma_P-\gamma_Q}$ and $q^* = \frac{\gamma_P}{\gamma_P-\gamma_Q} - \frac{\mu\gamma_P z_P}{\Omega}$ is not analytically tractable but the existence conditions correspond to $\lambda_P > 0$, $\lambda_Q > 0$ implying, by continuity considerations, that the fully mixed point is stable when it exists. The nature of the fixed point is inaccessible through analytic methods but it can either be a stable node or a stable oscillatory node according to the sign of the discriminant.

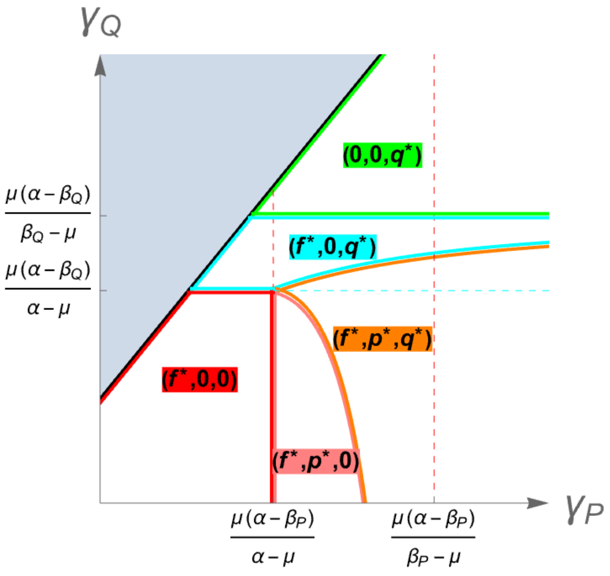


Figure 1: Phase portrait for plasmid compensation

A conventional explanation would assert that due to competitive exclusion the fully mixed solution should never be stable. An adaptive dynamics interpretation would only suggest that the linear invasion of the new plasmid would occur when introduced. Instead this full analysis, accompanied by numerical simulation results, indicates the existence of a complete coexistence regime for biologically relevant parameters.

### 3 Chromosome compensation model

The other salient case we consider is if another strain emerges which is assumed to carry a mutation that compensates the cost of the plasmid with a mutation on the chromosome. The new strain has a population fraction $c$, is assumed to be baring a plasmid and its properties are subscripted $C$, as opposed to $P$ for the original strain. The new strain has a differing plasmid conjugation rate $\gamma_C$ as compared to $\gamma_P$. The full system has dynamics

$$\begin{aligned}\frac{df}{dt} =& \alpha f(1-f-p-c) - \mu f - \gamma_P p f - \gamma_C c f \\ \frac{dp}{dt} =& \beta_P p(1-f-p-c) - \mu p + \gamma_P p f + \gamma_C c f \\ \frac{dc}{dt} =& \beta_C c(1-f-p-c) - \mu c.\end{aligned} \tag{6}$$

To match our assumptions we have chosen $\beta_P < \beta_C < \alpha$ (WLOG mathematically). Taking these inequalities forward we find the following Jacobian matrix and fixed points:

$$J(f,p,c) = \begin{bmatrix} \alpha(\Delta-f)-\mu-\gamma_P p-\gamma_C c & -\alpha f-\gamma_P f & -\alpha f-\gamma_C f \\ -\beta_P p+\gamma_P p+\gamma_C c & \beta_P(\Delta-p)-\mu+\gamma_P f & -\beta_P p+\gamma_C f \\ -\beta_C c & -\beta_C c & \beta_C(\Delta-c)-\mu \end{bmatrix} \tag{7}$$

with $\Delta = 1-f-p-c$.

- $(0,0,0)$, this fixed point – no bacteria – is always unstable due to our assumptions
- $(f^*,0,0)$, this fixed point, $f^* = 1-\dfrac{\mu}{\alpha}$ – no plasmid – is stable when $\gamma_P < \frac{\mu(\alpha-\beta_P)}{\alpha-\mu}$.
- $(0,p^*,0)$, this fixed point, $p^* = 1-\dfrac{\mu}{\beta_P}$ – original plasmid only – is always unstable
- $(f^*,p^*,0)$, a fixed point, $f^* = \frac{\alpha}{\gamma_Q z_Q} - \frac{\mu}{\gamma_Q}$, $p^* = \frac{\mu}{\gamma_Q} - \frac{\beta_Q}{\gamma_Q z_Q}$ – a mixed solution between frees and original plasmids.

- $(0, 0, c^*)$, this fixed point, $q^* = 1 - \frac{\mu}{\beta_C}$ – new plasmid dominates – is stable when $\gamma_C > \frac{\mu(\alpha - \beta_C)}{\beta_C - \mu}$
- $(f^*, p^*, c^*)$, a fixed point – a full mixed coexistence solution between all three strains.

This fully mixed fixed point is given by $f^* = \frac{(\beta_C - \beta_P)\gamma_C}{\beta_C \Sigma}(\beta_C - \mu z_C)$, $p^* = \frac{(\alpha - \beta_C)\gamma_C}{\beta_C \Sigma}(\beta_C - \mu z_C)$ and $c^* = \frac{(\alpha - \beta_C)\gamma_C}{\beta_C \Sigma}(\mu z_P - \beta_C)$ and we define $\Sigma = \Omega - \alpha(\gamma_P - \gamma_C)$ for convenience and $\Sigma$ and $z_{<.>}$ as before. This fixed point has two differing roles. When $\gamma_P$ is large and $\gamma_C$ small this is the stable fixed point which may or may not be oscillatory in character. A more applicable role is it acts as a boundary (it is a saddle in this situation) between two stable fixed points in the region marked ”3 fixed points”. This creates the possibility for dependence on initial conditions in this region and the inability of a chromosomal mutation to invade unless it does so with sufficient numbers.

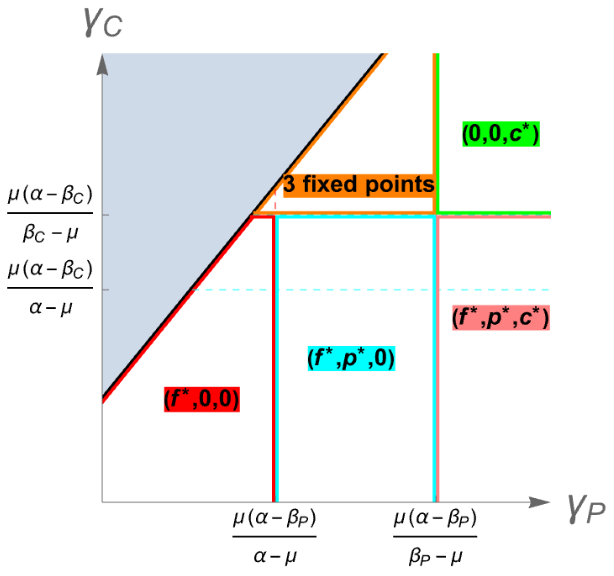


Figure 2: Phase portrait for chromosome compensation

#### 4 Selection

Selection can easily be imposed by adding a higher turnover term for any sensitive species. At its simplest, if we assume that the mechanism of resistance is identical, and is not affected by any subsequent compensation, this can be included as a linear loss term, proportional to $\eta$ on the plasmid free species.

For the basic model this gives the following equations:

$$\begin{aligned}\frac{df}{dt} =&\alpha f(1-f-p)-\mu f-\gamma pf-\eta f\\ \frac{dp}{dt} =&\beta p(1-f-p)-\mu p+\gamma pf,\end{aligned} \tag{8}$$

which shifts the free only fixed point to $1-\frac{\mu+\eta}{\alpha}$ and the mixed fixed point to $f_s^* = f^* - \eta\left(\frac{\beta}{z\gamma^2}\right)$,$p_s^* = p^* + \eta\left(\frac{\beta-\gamma}{z\gamma^2}\right)$, shifting the solution toward plasmid dominance, as expected.

For the plasmid compensation system the equations become

$$\begin{aligned}\frac{df}{dt} =&\alpha f(1-f-p-q)-\mu f-\gamma_P pf-\gamma_Q qf-\eta f\\ \frac{dp}{dt} =&\beta_P p(1-f-p-q)-\mu p+\gamma_P pf\\ \frac{dq}{dt} =&\beta_Q q(1-f-p-q)-\mu q+\gamma_Q qf,\end{aligned} \tag{9}$$

and analogously for the chromosome compensation system. In both cases the analysis can be computed exactly but the results do not change our interpretation. For small values of $\eta$ the findings are unchanged. A selection limit, $\eta^*$ exists (equal to the maximum of two value determined by the differing plasmids) where if selection is greater than both of these values then plasmid free states cannot exist and therefore the fittest plasmid solution always wins, which will be the compensated state, but it may take a long time due to relative absence of plasmid free target to facilitate spread. For the plasmid compensation

$$\eta_Q^* = \frac{z_Q\gamma_Q\mu - \beta_Q\gamma_Q}{\beta_Q} \tag{10}$$

whilst for the chromosome compensation

$$\eta_C^* = \frac{z_C\gamma_C\mu - \beta_C\gamma_C}{\beta_C} \tag{11}$$

which means as selection increases the compensated, plasmid containing bacteria will always dominate, as expected.
